# Functional analyses of the horizontally acquired Phytophaga glycoside hydrolase family 45 (GH45) proteins reveal distinct functional characteristics

**DOI:** 10.1101/489054

**Authors:** André Busch, Etienne G.J. Danchin, Yannick Pauchet

## Abstract

Cellulose, a major polysaccharide of the plant cell wall, consists of β-1,4-linked glucose moieties forming a molecular network recalcitrant to enzymatic breakdown. Although cellulose is potentially a rich source of energy, the ability to degrade it is rare in animals and was believed to be present only in cellulolytic microbes. Recently, it has become clear that some animals encode endogenous cellulases belonging to several glycoside hydrolase families (GHs), including GH45. GH45s are distributed patchily among the Metazoa and, in insects, are encoded only by the genomes of Phytophaga beetles. This study aims to understand both the enzymatic properties and the evolutionary history of GH45s in these beetles. To this end, we tested the enzymatic abilities of 37 GH45s derived from five species of Phytophaga beetles and learned that beetle-derived GH45s degrade three different substrates: amorphous cellulose, xyloglucan and glucomannan. Our phylogenetic and gene structure analyses indicate that at least one gene encoding a putative cellulolytic GH45 was present in the last common ancestor of the Phytophaga, and that GH45 xyloglucanases evolved several times independently in these beetles. The most closely related clade to Phytophaga GH45s contained fungal sequences, suggesting this GH family was acquired by horizontal gene transfer from fungi. Other than in insects, arthropod GH45s do not share a common origin and appear to have emerged at least three times independently.

## Introduction

The major source of energy for most organisms on earth is D-glucose. Through photosynthesis, plants have evolved the ability to biosynthesize organic D-glucose from inorganic carbon dioxide. A surplus of electromagnetic energy has provided plants with a nearly unlimited access to glucose, giving them a reservoir for storing energy as well as access to material for building structural components during plant growth. The plant cell wall (PCW) consists of several glucose-derived polysaccharides, which form a protective wall against biotic and abiotic stresses. Traditionally, three kinds of polysaccharides are used as structural elements in the PCW: cellulose, hemicellulose and pectin. While the latter two are characterized by a variety of differently organized heteropolysaccharides, cellulose is a homopolymer and consists of β-1,4-linked anhydroglucose units forming a straight-chain polysaccharide. Through hydrogen bonding, individual chains attach to each other and form a resilient (para)crystalline structure (Chang 1981). On surface areas, cellulose is believed to organize itself into a state of low crystallinity, referred to as an “amorphous” state (Ruel, et al. 2012). Depending on the developmental stage of the plant cell, the PCW is organized as follows: (i) the primary cell wall, which contains low amounts of crystalline cellulose (and surrounds plant cells in development) or (ii) the secondary cell wall, which comprises large amounts of crystalline cellulose (Cosgrove 2014). However, how native cellulose is organized in primary and secondary cell walls with regard to the ratio of amorphous to crystalline cellulose is still unclear (Saxena 2007; Knox 2008).

As the most abundant biopolymer on earth (Bayer, et al. 1998), cellulose represents an abundant energy supply for any organism which has the ability to exploit it. Curiously, cellulose degradation has evolved only in few branches of the tree of life. Until the end of the 20th century, cellulose degradation was only known to be performed by microorganisms such as plant pathogenic bacteria (Chambost J.P. 1987; Py, et al. 1991), saprophytic fungi (Schulein 1997) or mutualistic symbionts in insects and ruminants (Breznak and Brune 1994; Rincon, et al. 2001). However, in 1998, the first endogenous cellulases of animal origin were identified in cyst nematodes found in parasitic plants (Smant, et al. 1998) and termites (Watanabe, et al. 1998). Several other independent discoveries of cellulases in a variety of Metazoa followed, and to date endogenous cellulases encompass the phyla Arthropoda, Mollusca and Nematoda (Girard and Jouanin 1999; Kikuchi, et al. 2004; Sakamoto and Toyohara 2009; Pauchet, et al. 2010; Faddeeva-Vakhrusheva, et al. 2016).

Cellulases are conventionally classified according to their mode of action. Endo-β-1,4,-glucanases (EC 3.2.1.4) break down cellulose by releasing randomly sized cellulose fragments and are known to act only on amorphous cellulose. Cellobiohydrolases (exo-β-1,4,-glucanases; EC 3.2.1.91) degrade cellulose from its terminal regions by releasing cellobiose and occasionally cellotriose. In microbes, cellobiohydrolases were shown to degrade amorphous as well as crystalline cellulose (Takahashi, et al. 2010). Finally, cellobiosidases (β-glucosidases; EC 3.2.1.21) accept the released cellobiose as substrate and convert it into glucose. All three types of cellulases act synergistically and are necessary to degrade the cellulosic network efficiently (Kostylev and Wilson 2012).

Cellulases are distributed into 14 of the 156 currently described families of glycoside hydrolases (GHs), according to the carbohydrate-active enzyme (CAZy) database (www.cazy.org, (Lombard, et al. 2014). Assignment to different GH families is based on sequence similarities. The best-described cellulolytic GH families encompass GH5 (Aspeborg, et al. 2012) and GH9 (Watanabe and Tokuda 2010). Together with GH45s, GH5s and GH9s are found to be encoded by the genome of some insects (Keeling, et al. 2013; Pauchet, Saski, et al. 2014; Vega, et al. 2015; McKenna, et al. 2016; Schoville, et al. 2018). Based on our previous work, GH45s are commonly distributed in the Phytophaga clade of beetles (Marvaldi, et al. 2009), which encompasses the superfamilies Chrysomeloidea (leaf beetles and longhorned beetles) and Curculionoidea (weevils and bark beetles) (Pauchet, et al. 2010; Kirsch, et al. 2012; Pauchet, Kirsch, et al. 2014). The first GH45 that was functionally characterized in a beetle originated from *Apriona germari* (Chrysomeloidea: Cerambycidae: Lamiinae), which had the ability to degrade amorphous cellulose (Lee, et al. 2004). Until recently, GH45s in beetles have been functionally characterized in only a few Chrysomeloidea species, mostly Cerambycidae (Chang, et al. 2012; Pauchet, Kirsch, et al. 2014; Mei, et al. 2015) and *Diabrotica virgifera virgifera* (Chrysomelidae: Galerucinae) (Valencia, et al. 2013), and another two from our previous study in *Gastrophysa viridula* (Chrysomelidae: Chrysomelinae) (Busch, et al. 2018). Although GH45 sequences have been identified in Curculionoidea beetles (Pauchet, et al. 2010; Keeling, et al. 2013; Vega, et al. 2015), to date none has ever been functionally characterized.

Interestingly, GH45s are not only found in multicellular organisms but are widely encoded by microbes (Sheppard, et al. 1994; Davies, et al. 1995; DeBoy, et al. 2008; O’Connor, et al. 2014). The distribution of this gene family in the Metazoa appears to be patchy and has so far been recorded only in a few species within the phyla Mollusca (Xu, et al. 2002; Sakamoto and Toyohara 2009; Rahman, et al. 2014), Nematoda (Kikuchi, et al. 2004; Palomares-Rius, et al. 2014); GH45s have also been recorded in Arthropoda (Song, et al. 2017). If GH45s had evolved in the last common ancestor (LCA) of the Metazoa and subsequently been inherited by their descendants, we would expect the patchy distribution of GH45 genes observed within the Metazoa to be due to multiple independent losses. If this hypothesis were true, phylogenetic analyses would recover metazoan GH45s as a single monophyletic clade. However, two previous studies focusing on the evolutionary origin of GH45s in nematodes and mollusks have suggested instead that GH45s were acquired from a fungal donor by horizontal gene transfer (HGT) (Kikuchi, et al. 2004; Sakamoto and Toyohara 2009). The first attempt to clarify the evolutionary history of GH45s in beetles also proposed an HGT from a fungal source but was unable to reach definite conclusions (Calderon-Cortes, et al. 2010) because of the low number of sequences used for the phylogenetic analysis. A more comprehensive approach followed in 2014 (Eyun, et al.), which included more GH45 sequences. Still, the variety of GH45 sequences in the latter study was poor, resulting in a similarly elusive outcome. Thus, the evolutionary history of GH45s appears to be complex, and their inheritance in beetles remains enigmatic.

Therefore, the major aim of our study was to trace the evolutionary origin of the GH45 family within the Phytophaga and to analyze how the function of the corresponding proteins evolved in this large group of beetles. Based on previous research on the ancestral origin of GH45s (Kikuchi, et al. 2004; Sakamoto and Toyohara 2009), we hypothesize that an HGT event occurred at one or more stages of the evolution of the Phytophaga. Additionally, we analyzed other Arthropods, including Oribatida and Collembola, as well as several non-arthropod species, including Nematoda, Tardigrada and Rotifera. In this study, we combined functional and phylogenetic analyses to unravel the origin and evolution of the GH45 family in Phytophaga beetles. We first functionally characterized 37 GH45s from five beetle species -- four beetles of the Chrysomelidae (leaf beetles) and a beetle of the Curculionidae (weevils) -- to determine whether these GH45s harbor cellulase activity, and whether they may have evolved other functions. We then combined these functional data with amino acid alignments of the GH45 catalytic sites to pinpoint amino acid substitutions which might lead to substrate shifts. Finally, we performed phylogenetic analyses to ask (i) how many GH45 genes were present in the LCA of the Phytophaga and (ii) whether this gene family is ancestral in the Metazoa or, instead, acquired by HGT. The aim of our study was to provide the first comprehensive overview regarding the evolution in beetles of the GH45 family and to assess the role of these genes in the evolution of herbivory.

## Results

### Functional analyses of the Phytophaga GH45 proteins reveal distinct enzymatic characteristics

Our previous transcriptome analyses (Pauchet, et al. 2010; Kirsch, et al. 2012; Eyun, et al. 2014) revealed a set of endogenous GH45 genes distributed within several beetles of the superfamilies Chrysomeloidea and Curculionoidea. We investigated the product of GH45 genes from four beetle species belonging to the family Chrysomelidae, namely, *Chrysomela tremula* (CTR), *Phaedon cochleariae* (PCO), *Leptinotarsa decemlineata* (LDE) and *Diabrotica virgifera virigfera* (DVI), and from one species belonging to the family Curculionidae, the rice weevil *Sitophilus oryzae* (SOR), for a total of 33 GH45 sequences (Table S1). By re-examining the corresponding transcriptomes as well as the recent draft genome of *L. decemlineata* (Schoville, et al. 2018), we identified four extra GH45 sequences (Table S1). The resulting 37 GH45s were successfully expressed in *Sf*9 insect cells (Fig. 1A). All GH45s had an apparent molecular weight of ~35 kDa (Fig. 1A). The increase in molecular size compared to the expected size (~25 kDa) was likely due to post-translational N-glycosylations as well as to the artificially added V5/(His)_6_-tag.

**Figure 1.**
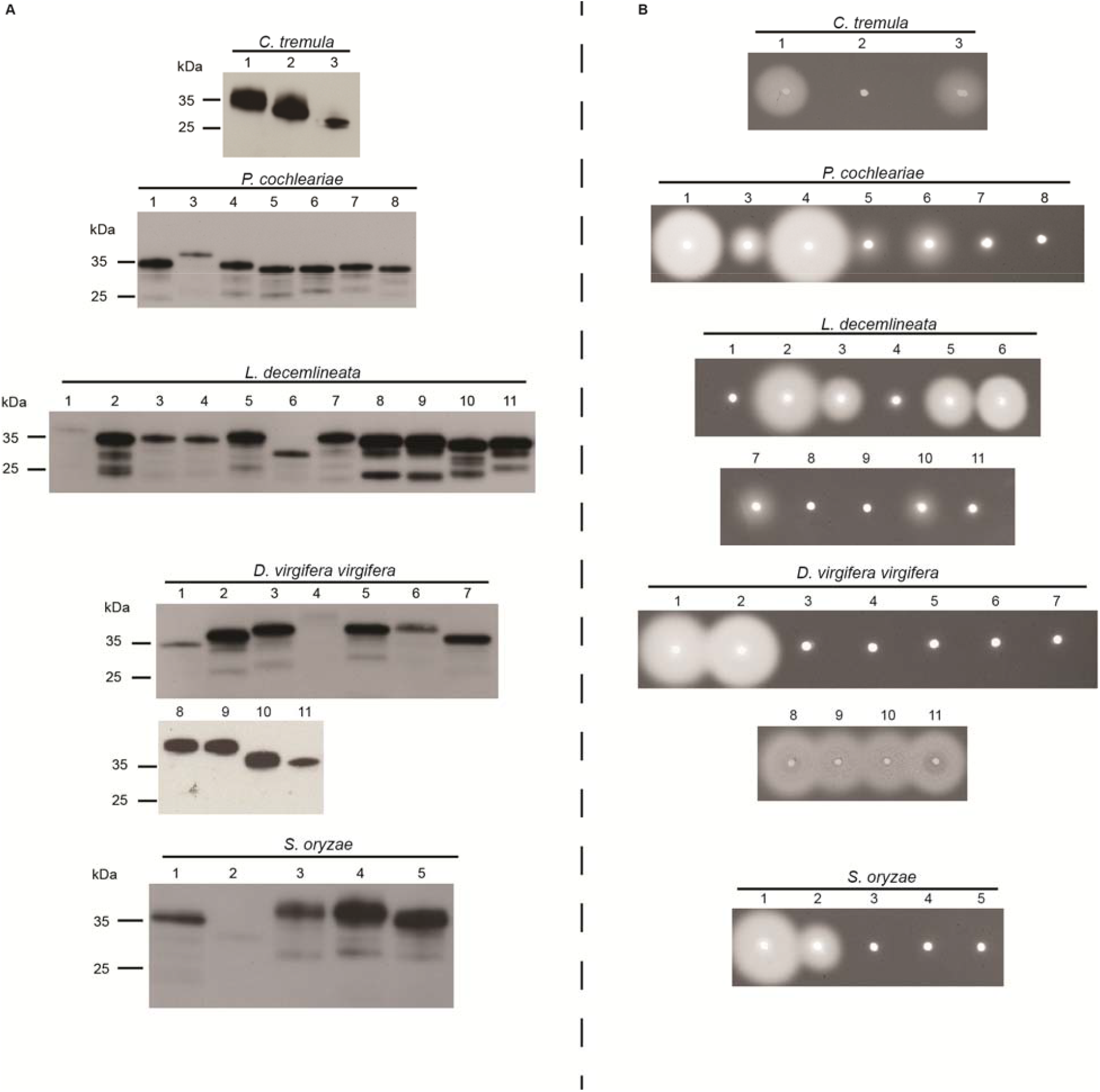
Western blot and CMC-based agarose-diffusion assay of target GH45 proteins. A) Western blot of target recombinant enzymes expressed in frame with a V5/(His)_6_ after heterologous expression in insect *Sf*9 cells. After 72 h, crude culture medium of transfected cells was harvested and analyzed by Western blotting using an anti-V5 HRP-coupled antibody. B) Crude culture medium of transfected cells was applied to an agarose-diffusion assay containing 0.1 % CMC. Activity halos were revealed after 16 h incubation at 40 °C using Congo red. Numbers above Western blot and agarose-diffusion assays correspond to the respective species of GH45s depicted in supplementary Table S1.

To explore the cellulolytic capabilities of these proteins, we first applied crude *Sf*9 culture medium containing individual recombinant GH45s to agarose plates supplemented with 0.1 % carboxymethyl cellulose (CMC) (Fig. 1B). Activity halos were visible for at least two GH45s per target species. The intensity of the observed activity halos varied from large clearing zones (for example, PCO4 or LDE2) to small or medium ones (such as PCO3 or LDE7). These differences were likely due to the catalytic efficiency of each individual GH45 as well as to the concentration of the crude protein extracts we used. Each clearing zone, independent of its intensity and size, indicated endo-β-1,4-glucanase activity.

To further assess enzymatic characteristics of these GH45s, we performed assays with a variety of plant cell wall-derived polysaccharides as substrates and analyzed the resulting breakdown products on thin layer chromatography (TLC) (Table 1; Fig. S1 to S5). We were able to confirm the cellulolytic activity initially observed on CMC agar plates (CTR1 and CTR3, PCO1, PCO3, PCO4 and PCO6, LDE2, LDE3, LDE5, and LDE6, DVI1, DVI2 and DVI8-11, SOR1 and SOR2). Each of these enzymes was able to break down CMC, regenerated amorphous cellulose (RAC) and cellulose oligomers. Interestingly, LDE10 did not show activity against cellulosic polymers but preferentially degraded cellopentaose and cellohexaose (Fig. S3). Similarly, but with much weaker efficiency, PCO5 degraded cellohexaose (Fig. S2). Together with the plate assays, our TLC analyses clearly demonstrated that beetle-derived GH45s processed cellulosic substrates using an endo-active mechanism, which suggests that these enzymes are endo-β-1,4-glucanases. Several cellulolytically active GH45s derived from the four leaf beetle species displayed additional activity towards the hemicellulose glucomannan, for example, CTR1, PCO3 and LDE2 (Table 1; Figs. S1 to S3). LDE7 exhibited the highest enzymatic activity against glucomannan, whereas its activity against amorphous cellulose substrates could be visualized only by plate assay.

**Table 1:**
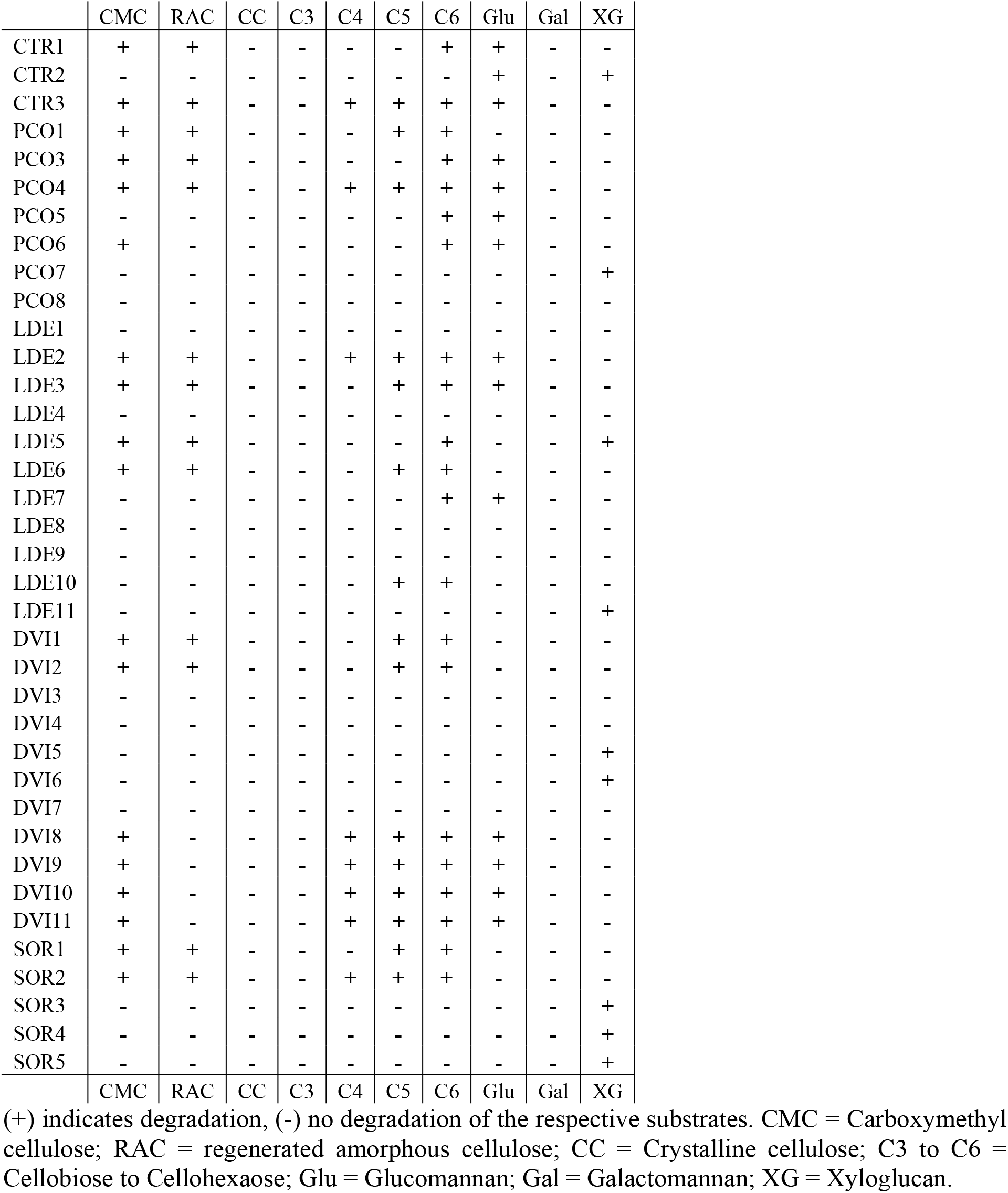
GH45 enzymatic activity observed on TLC.

Interestingly, TLC allowed us to detect several enzymes (CTR2, PCO7, LDE5, LDE11, DVI5, DVI6 and SOR3 to SOR5) which were able to degrade xyloglucan instead of cellulose (Table 1; Figs. S1 to S5). The size of the resulting breakdown products seemed to correlate with heptamers and octamers, indicating that these proteins were endo-β-1,4-xyloglucanases; however, the actual size of the resulting breakdown products is difficult to assess because some glucose moieties that make up the backbone of xyloglucan are substituted with xylose residues. LDE5 displayed activity towards amorphous cellulose substrates (Figs. 1B and S3), and is thus, according to our data, the only example of a beetle-derived GH45 able to degrade xyloglucan as well as amorphous cellulose. Additionally, each of the 37 GH45s was tested against xylan and no activity was detected (data not shown).

The Chrysomelid-derived PCO8, LDE1, LDE4, LDE8, LDE9, DVI3, DVI4 and DVI7 did not exhibit activity towards any of the substrates used in this study. We wondered whether substitutions of catalytically important residues may have caused their apparent loss of activity. To test this hypothesis, we performed an amino acid alignment of target beetle GH45 sequences, including a fungal GH45 sequence, as a reference for which the structure has been resolved (Davies, et al. 1995); then we screened for amino acid substitutions and compared these to the reference fungal sequence (Fig. 2). According to Davies et al. (1995), both the proton donor (catalytic acid) and the acceptor (catalytic base) of the catalytic dyad should be aspartates (Asp10 and Asp121). In LDE9, the catalytic base was substituted for an asparagine, whereas in DVI4, the catalytic acid was substituted for a valine (Fig. 2). In both cases, the loss of the carboxyl unit of the functional group likely caused the proteins to lose the catalytic activity. No substitution event of the catalytic residues was observed for PCO8, LDE1, LDE4, LDE8, DVI3, and DVI7. Thus, we decided to investigate several conserved sites known to affect the enzymatic activity of GH45s (Davies, et al. 1995). In addition, we investigated three other sites crucial for enzymatic activity: (i) a proposed stabilizing aspartate (Asp114), (ii) a conserved alanine (Ala74) and (iii) a highly conserved tyrosine (Tyr8) (Fig. 2). Apart from two substitution events of Tyr8 in LDE9 and LDE3, this amino acid remained conserved in all other beetle GH45 sequences. LDE9 already possessed a mutation in its catalytic acid, which was likely responsible for the lack of activity. In LDE3, a substitution from Tyr8 to Phe8 did not significantly impact the catalytic abilities of this protein, likely because the side-chains of both amino acids are highly similar and differ only in a single hydroxyl group. When examining the proposed stabilizing site Asp114, we observed several amino acid substitutions that correlated with a loss of activity in PCO8, DVI3, DVI4, LDE1 and LDE8. Amino acid changes at the Asp114 position were also observed in PCO3 and CTR1, but were not correlated with a loss of enzymatic activity. Since LDE4 and DVI7 appeared to have no mutation in Asp114, we screened the Ala74 residue for substitutions; the amino acid exchange we observed, from alanine to glycine in both cases, may have caused the loss of activity in these two proteins. Altogether, amino acid substitutions at important sites could be detected in some apparently inactive GH45s, but not in all of them. It may be that the proteins for which we did not find amino acid substitutions are still active enzymes, and we have just not yet found the right substrate; alternatively, we have not yet checked all the amino acid positions, some of which could also be crucial for catalysis.

**Figure 2.**
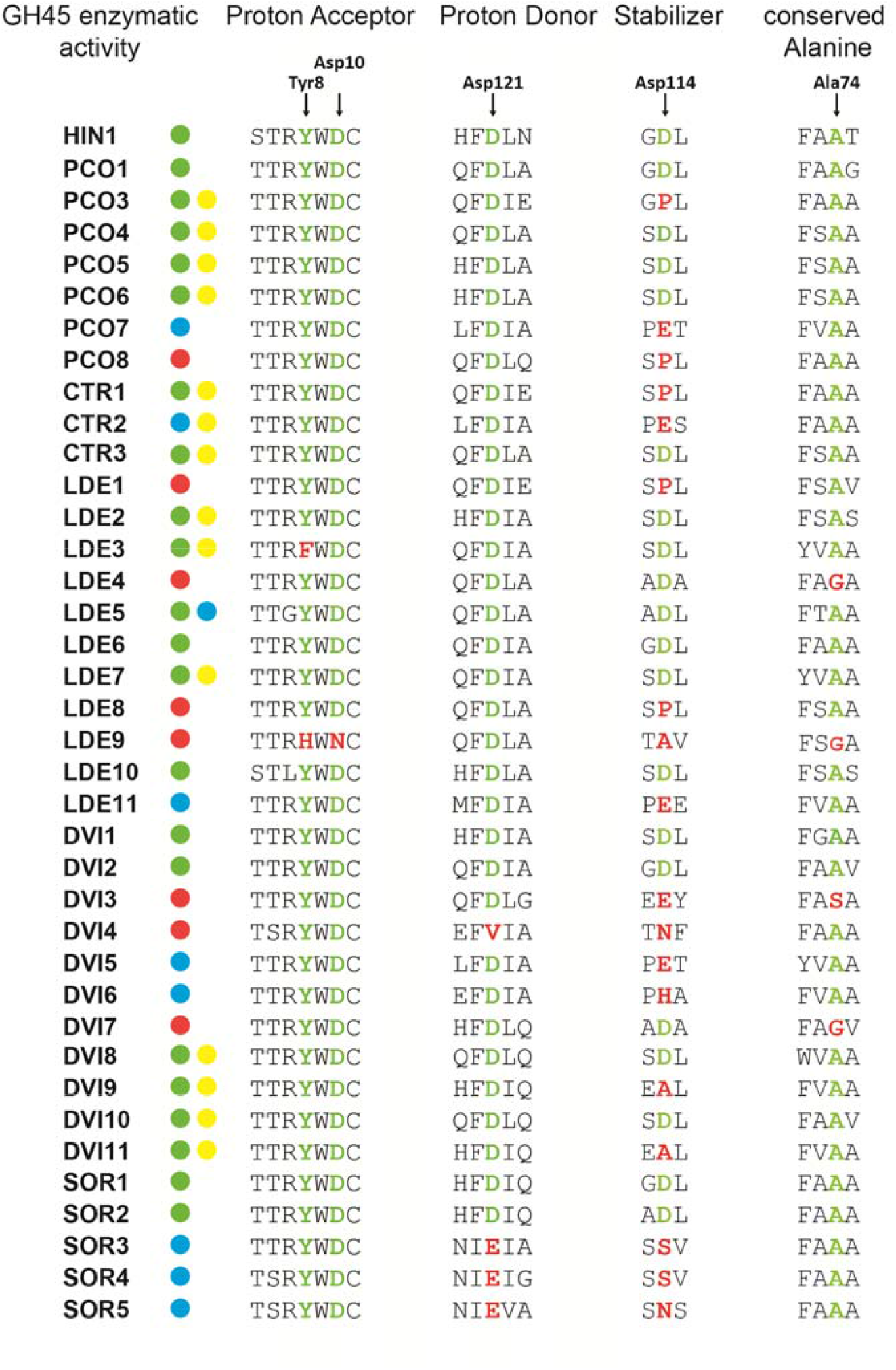
GH45 amino acid alignment of the catalytic residues. We used a GH45 sequence of *Humicola insulens* (HIN1) as a reference sequence (Accession: 2ENG_A) (Davies, et al. 1995). According to HIN1, we chose to investigate the catalytic residues (ASP10 and ASP121) as well as a conserved tyrosine (TYR8) of the catalytic binding site, a crucial substrate-stabilizing amino acid (ASP114) and an essential conserved alanine (ALA74). Arrows indicate amino acid residues under investigation. If the respective amino acid residue is highlighted in green, it is retained in comparison to the reference sequence; otherwise it is highlighted in red. GH45 enzymatic activity was color-coded based on the respective substrate specificity (green dots = endo-β-1,4-glucanase, blue dots = endo-β-1,4-xyloglucanase, yellow dots = (gluco)mannanase, red dots = putatively non-activity).

Interestingly, all Chrysomelid GH45 xyloglucanases (except LDE5 and DVI6), including *G. viridula* GVI1 from our previous study, (Busch, et al. 2018), displayed a substitution from aspartate to glutamate at the stabilizing site (Asp114) (Fig. 2). As glutamate differs from aspartate only by an additional methyl group within its side chain, we believe that this exchange may have contributed to the substrate shift. Interestingly, and in contrast to Chrysomelidae-derived GH45 xyloglucanases, we found that GH45 xyloglucanases from the Curculionidae *S. oryzae* (SOR3 to SOR5) had a substitution from aspartate to glutamate in the proton donor residue (Asp121). We also believe that, in *S. oryzae*, this particular substitution may have contributed to the preference for xyloglucan over cellulose as a substrate.

In summary, we demonstrated that each species investigated encoded at least two cellulolytic GH45s that are able to degrade amorphous cellulose. We also demonstrated that at least one GH45 per species possessed the ability to degrade only xyloglucan. Interestingly, several GH45s did not show activity on any of the substrates we tested, suggesting that they have become pseudo-enzymes or are active on substrates not tested here.

### Phylogenetic analyses reveal multiple origins of GH45 genes during the evolution of Metazoa

For further insight into the evolutionary history of beetle-derived GH45 genes, we used phylogenetic analyses to reconstruct their evolutionary history. To achieve this goal, we collected amino acid sequences of GH45s available as of February 2018, including those from the CAZy database (http://www.cazy.org) (Lombard, et al. 2014) as well as from several transcriptome datasets accessible at NCBI Genbank. Interestingly, we realized that the presence of GH45 genes in arthropods was not restricted to Phytophaga beetles: these genes were also distributed in transcriptomes/genomes of species of springtails (Hexapoda: Collembola) and of species of Oribatida mites (Arthropoda: Chelicerata) (Table S3). In addition, we identified GH45 sequences in two other groups of Metazoa, namely, tardigrades and rotifers. Notably, our homology search did not retrieve any mollusk-derived GH45s; given the distant relationship of these GH45s to any of those we investigated, this absence is not surprising. The patchy distribution of GH45 genes throughout the arthropods and, more widely, the Metazoa, could be due to either the presence of GH45 genes in a common ancestor, followed by multiple gene losses, or from multiple independent acquisitions from foreign sources (i.e., HGT). To test these hypotheses, we collected a diverse set of GH45 sequences of microbial and metazoan origins resulting in 264 sequences (Table S3). Subsequently, redundancy at 90% identity level between sequences was eliminated, resulting in 201 non-redundant GH45 sequences. According to both Bayesian and maximum likelihood phylogenies (Fig. 3, S6 andS7), the arthropod-derived GH45 sequences were not monophyletic but globally fell into three separate groups. One highly supported monophyletic clade (posterior probability (PP) =1.0, bootstrap =85) grouped all the Phytophaga beetle GH45 sequences. This clade was most closely related to a group of Saccharomycetales fungi (PP=0.88, bootstrap=44). Then this clade branched to yet another group of Saccharomycetales Fungi (PP= 1.0, bootstrap=56). A second monophyletic clade grouped all the Oribatida mites GH45 sequences, although with moderate support on the branch (PP=0.72, bootstrap=3). Finally, a third clade (PP=0.96, bootstrap=14) grouped all the GH45 sequences from Collembola. This last group was not monophyletic: a bacterial GH45 sequence was interspersed within this clade and separated the Collembola GH45 sequences into two subgroups. In addition to the arthropods, the nematode GH45 sequences formed a highly supported monophyletic clade (PP= 1.0, bootstrap=84) which was connected to a clade of fungal-derived sequences. This connection was highly supported (PP= 0.93, bootstrap=69). The two other groups of Metazoa (tardigrades and rotifers) were located in a separate clade with species of Neocallimastigaceae fungi (Chytrids), (PP=0.94, bootstrap=14). Overall, this analysis showed that neither arthropods nor, more generally, metazoan GH45 sequences, originated from a common ancestor, as they were scattered in multiple separate clades rather than forming a monophyletic metazoan clade.

**Figure 3.**
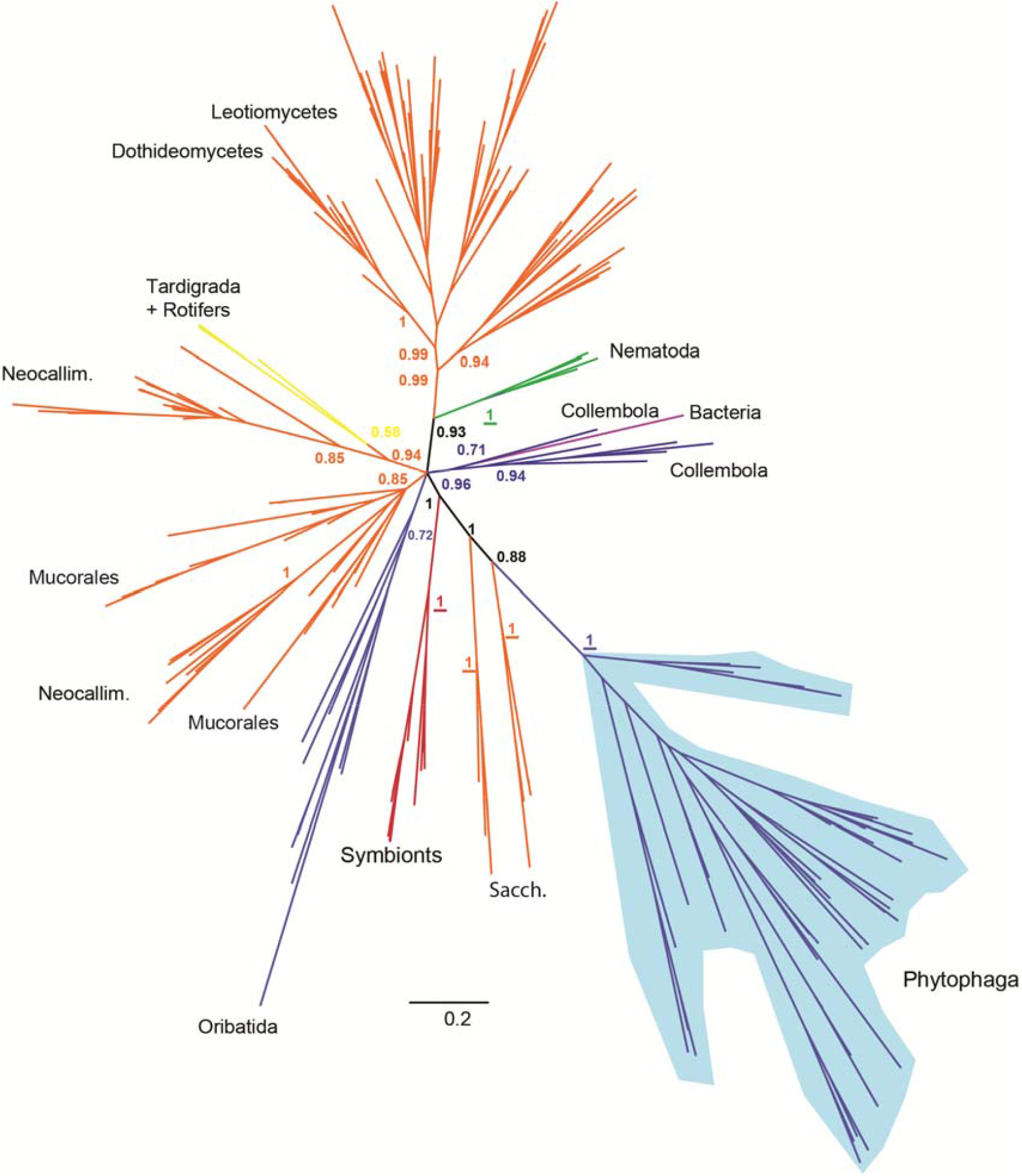
Global phylogeny encompassing GH45 proteins from various taxa. Bayesian-based phylogenetic analysis of GH45 sequences. 264 GH45 sequences of microbial and metazoan origin were initially collected (see Methods), and their redundancy was eliminated at 90 % sequence similarity, resulting in a total of 201 sequences. Posterior probability values are given at crucial nodes. If values are depicted in bold, the same branch appeared in the corresponding maximum likelihood analysis (see Figs. S6 and S7). If underlined, the maximum likelihood node was highly supported (bootstrap values > 75). Detailed sequence descriptions including accession numbers are given in Table S3. Hexapoda are represented in blue, fungi in orange, protists in red, bacteria in purple, Nematoda in green and other Metazoa in yellow. Sacch. = Saccharomycetales fungi; Neocallim. = Neocallimastigaceae fungi.

In summary, our phylogenetic analyses illustrated that the evolutionary history of GH45s in the Metazoa was complex and pointed to the possibility that this gene family evolved several times independently in multicellular organisms. More specifically, our analyses suggested that this gene family had evolved at least three times independently in arthropods. Finally, our data pointed toward an acquisition of GH45 genes by the LCA of Phytophaga beetles -- presumably through an HGT event -- from a fungal donor.

### The structure of GH45 genes in Phytophaga beetles supports a single origin before the split of the Chrysomeloidea and Curculionoidea

The monophyly of the Phytophaga-derived GH45s in the above phylogenetic analyses suggests a common ancestral origin in this clade of beetles. If the presence of a GH45 in the Phytophaga beetles had resulted from a single acquisition in their LCA, we hypothesized that the GH45 genes present in current species of leaf beetles, longhorned beetles and weevils would share a common exon/intron structure. To test this hypothesis, we mined the publicly available genomes of three species of Curculionidae, including *H. hampei* (Vega, et al. 2015), *D. ponderosae* (Keeling, et al. 2013) and *S. oryzae* (unpublished), as well as the genomes of the Chrysomelidae *L. decemlineata* (Schoville, et al. 2018) and of the Cerambycidae *A. glabripennis* (McKenna, et al. 2016). We were able to retrieve the genomic sequence corresponding to each of the GH45 genes present in these beetle species, with the exception of DPO9, which we did not find at all, and SOR3 and SOR4, which we were able to retrieve only as partial genomic sequences. Our results showed that the number of introns varied between the different species (Fig. S8). In *L. decemlineata* (representing Chrysomelidae), we identified a single intron in each of the GH45 genes (except for LDE11, which had two). For *A. glabripennis* (representing Cerambycidae), we found two introns in each of the two GH45 genes. In *H. hampei, D. ponderosae* and *S. oryzae* (all representing Curculionidae), the number of introns ranged from three to five. Interestingly, all GH45 genes in these five species possessed an intron placed within the part of the sequence encoding the predicted signal peptide. Apart from DPO7 and DPO8, these introns were all in phase one. This gene structure of Chrysomelid- and Curculionid-derived GH45 genes correlated well with our previous study investigating the gene structure of PCW-degrading enzymes, including GH45 genes, in the leaf beetle *Chrysomela tremula* (Pauchet, Saski, et al. 2014). The conservation of the phase and the position of this intron indicated that the LCA of the Phytophaga likely possessed a single GH45 gene having a phase one intron located in a part of the sequence encoding a putative signal peptide. To assess whether that particular intron is also present in the most closely related fungal species, we blasted the genomes of Saccharomycetaceae and Neocallimastigaceae fungi (NCBI, whole-shotgun genome database) using the protein sequence of GH45-1 of *C. tremula*. We did not detect any introns in fungal GH45 sequences (data not shown), suggesting that the proposed intron was acquired after the putative HGT event. The diversity of the overall intron-exon structure in phytophagous beetles likely resulted from subsequent and independent intron acquisition. In summary, and together with the monophyly of beetle-derived GH45s (Fig. 3), our analysis highly supports a common ancestral origin of beetle GH45.

### Evolution of the GH45 family after the initial split of Chrysomeloidea and Curculionoidea

We mined publicly available transcriptome and genome datasets of Phytophaga beetles (Table S4) and collected as many GH45 sequences as possible. We curated a total of 266 GH45 sequences belonging to 42 species of Phytophaga beetles. After amino acid alignment, we decided to exclude 60 partial GH45 sequences from our phylogenetic analysis because these were too short. We performed a “whole Phytophaga” phylogenetic analysis on the remaining 206 curated GH45 sequences using maximum likelihood (Fig. 4). Because most of the deeper nodes were poorly supported, we decided to collapse branches having a bootstrap support below 50. Our phylogenetic analysis indicated that no orthologous genes were found between species of Chrysomeloidea and Curculionoidea (Fig. 4). The only exception to this rule was found in a clade containing the xyloglucanases from the Chrysomelidae (clade m) and from the Curculionidae (clade n), which cluster together with a bootstrap support of 73 Proceeding cautiously, because the substrate switch from amorphous cellulose to xyloglucan seemed to be due to different amino acid substitutions at catalytically important sites between Chrysomelidae-derived xyloglucanases and Curculionidae-derived ones (Fig. 2), we hypothesize a single common ancestry of cellulolytic GH45s; in contrast, xyloglucanase activity likely arose through convergent evolution at list twice within the Phytophaga clade of beetles.

**Figure 4.**
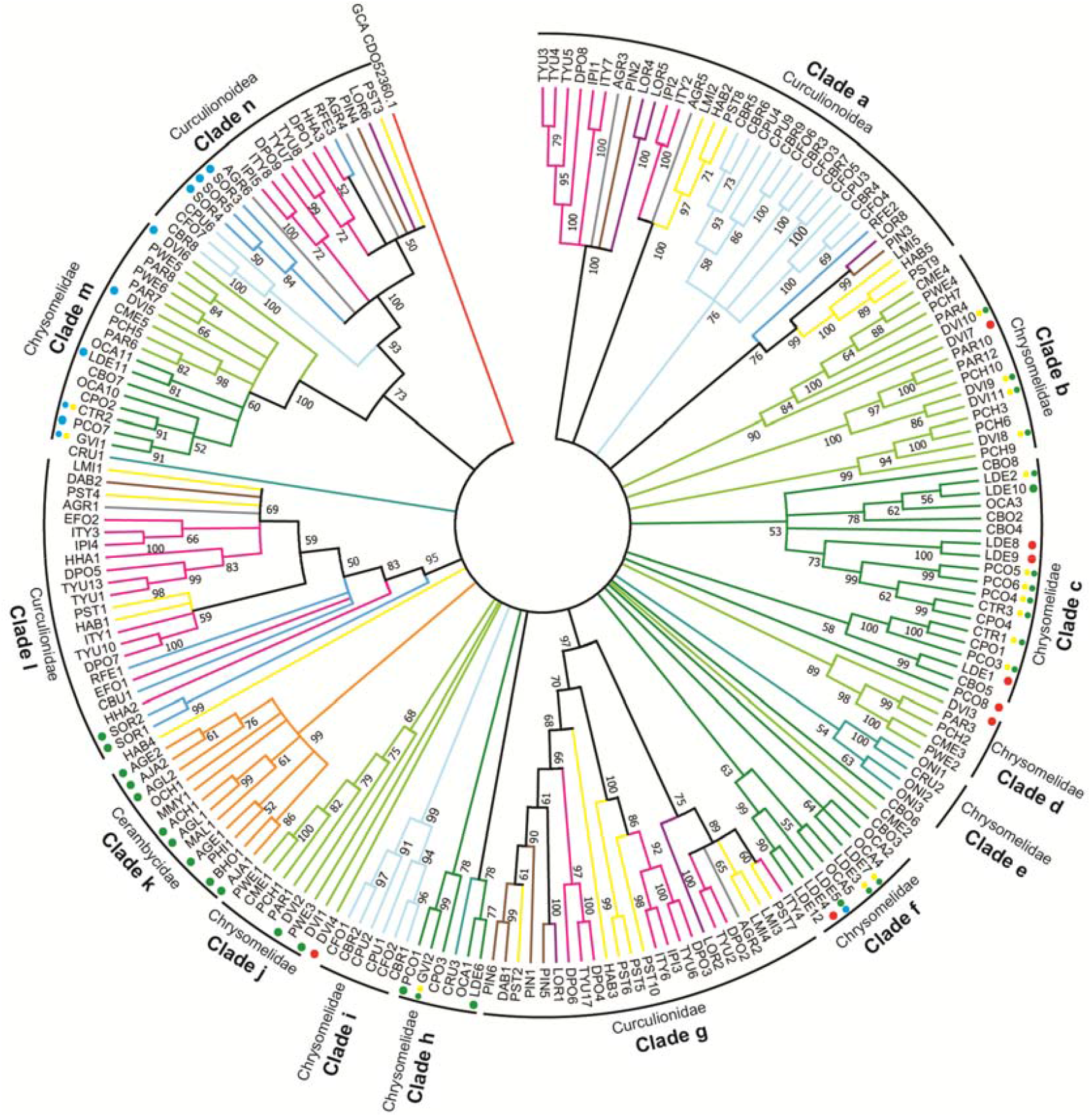
Phylogenetic relationships of Phytophaga-derived GH45s. A maximum-likelihood-inferred phylogeny of the predicted amino acid sequences of beetle-derived GH45s. Bootstrap values are indicated at corresponding nodes. The tree was collapsed at nodes below a bootstrap value of 50. Information on sequences and their accession number are given in Table S3. Dots indicate GH45s characterized to date and are color-coded based on their activity: green = endo-β-1,4-glucanases; blue = endo-β-1,4-xyloglucanases; yellow = (gluco)mannanases; red = no activity detected. Color coding in reference to the respective subfamily of Curculionoidea: pink = Scolytinae (Curculionidae); brown = Entiminae (Curculionidae); purple = Cyclominae (Curculionidae); gray = Curculioninae (Curculionidae); yellow = Molytinae (Curculionidae); light blue = Brentinae (Brentidae); dark blue = Dryophthorinae (Curculionidae). Color coding in reference to the respective subfamily of Chrysomeloidea: dark green = Chrysomelinae (Chrysomelidae); light green = Galerucinae (Chrysomelidae); orange = Lamiinae (Cerambycidae); cyan = Cassidinae (Chrysomelidae). GH45 enzymatic activity was color-coded based on the respective substrate specificity (green dots = endo-β-1,4-glucanase, blue dots = endo-β-1,4-xyloglucanase, yellow dots = (gluco)mannanase, red dots = putatively non-activity).

Clade n comprised Brentidae- and Curculionidae-derived GH45s including SOR3 to SOR5 (Fig. 4). According to our functional data, SOR3 to SOR5 act as xyloglucanases, suggesting that other GH45 proteins present in clade n may have also evolved to degrade xyloglucan. To support this hypothesis, we compared the catalytic residues of SOR3-5 to those of the other Curculionidae and Brentidae-derived GH45 sequences from clade n (Fig. S9). We detected substitutions from an aspartate to a glutamate at Asp121 in all Curculionidae-derived GH45 sequences of this clade but not in the Brentidae-derived sequences, suggesting that Curculionidae-derived GH45s of this clade were likely to possess xyloglucanase activity. Functional analyses of the Brentidae-derived sequences present in clade n will be needed to determine whether these proteins are also xyloglucanases or whether they fulfill another function.

The second major cluster encompassed GH45s of clade l, with a bootstrap support of 95, and contained only Curculionidae-derived sequences (Fig. 4). Within this clade, we found SOR1 and SOR2, which are, according to our functional data, endo-active cellulases. Their presence in clade l implies that other GH45s of this cluster exhibit potential endo-cellulolytic activity. To further support this hypothesis, we again investigated amino acid residues of the catalytic site by comparing SOR1 and SOR2 to other GH45 sequences in this clade (Fig. S9). We did not find crucial substitutions in any of the investigated sites, implying that all GH45 proteins of this clade may have retained endo-β-1,4-glucanase activity. GH45 sequences present in the other Curculionidae and Brentidae-specific clades did not harbor any amino acid substitutions which could impair their catalytic properties, and they may all possess the ability to break down amorphous cellulose. More functional analyses will be necessary to assess the function of these proteins.

Regarding Chrysomeloidea-derived sequences, a highly supported clade (Fig. 4, clade m) contained GH45 sequences of two subfamilies of Chrysomelidae, namely, Chrysomelinae and Galerucinae. Our functional analyses revealed that this clade contained GH45 proteins possessing xyloglucanase activity, including DVI5 and DVI6 from *D. vir. virgifera*. The remaining Galerucinae-derived GH45s, such as those from the Alticines *Phyllotreta armoraciae* and *Psylliodes chrysocephala*, present in clade m have yet to be functionally characterized. When the catalytic residues of active xyloglucanases from our study were compared to the uncharacterized GH45 sequences present in clade m, we observed that at least PAR6 and PCH5 of the Galerucinae and OCA10 and CPO2 of the Chrysomelinae had congruent substitutions (ASP114 > Glu114), which likely enabled those proteins to also degrade xyloglucan (Fig. S10). Therefore, it is highly likely that the LCA of the Chrysomelinae and the Galerucinae possessed at least two GH45 proteins, an endo-acting cellulase and a xyloglucanase.

Clade k consisted solely of Lamiinae-derived sequences and in fact encompassed all Cerambycidae-derived GH45s identified to date (Fig. 4). Several of those GH45s had been previously functionally characterized as cellulases including AJA1 and AJA2 (Pauchet, Kirsch, et al. 2014), AGE1 and AGE2 (Lee, et al. 2004; Lee, et al. 2005), AGL1 and AGL2 (McKenna, et al. 2016), ACH1 (Chang, et al. 2012) and BHO1 (Mei, et al. 2015). Investigating their catalytic residues revealed no critical substitutions (Fig. S10), indicating that each yet-uncharacterized Cerambycidae GH45s from clade k (i.e. OCH1, MMY1, PHI and MAL1) may also possess cellulolytic activity.

In summary, our focus on Phytophaga-derived GH45s with regards to enzymatic characterization and ancestral origin allowed us to postulate that at least one GH45 protein was present in the LCA of the Phytophaga beetles and that this GH45 protein likely possessed cellulolytic activity. After the split between Chrysomeloidea and Curculionoidea, the GH45 gene family evolved through gene duplications at the family, subfamily and even genus/species level. Finally, according to our data, the ability of these beetles to break down xyloglucan, one of the major components of the primary plant cell wall, happened at least twice, once in the LCA of the Chrysomelinae and Galerucinae and once in the LCA of the Curculionidae.

## Discussion

In our previous research, we found that several beetles of the Phytophaga encoded a diverse set of GH45 putative cellulases (Pauchet, et al. 2010). Here we demonstrated that in each of the five Phytophaga beetles investigated, at least two of these GH45s possess cellulolytic activity. This discovery is in accordance with other previously described GH45 proteins from Insecta (Pauchet, Kirsch, et al. 2014), Nematoda (Kikuchi, et al. 2004), Mollusca (Rahman, et al. 2014), Rotifera (Szydlowski, et al. 2015) and microbes (Mcgavin and Forsberg 1988).

Surprisingly, several GH45 proteins were able to degrade glucomannan in addition to cellulose. We hypothesize that GH45 bi-functionalization may have occurred as a result of the chemical similarities between cellulose and glucomannan. Glucomannan is a straight chain polymer consisting of unevenly distributed glucose and mannose moieties. GH45 cellulases could recognize two adjoining glucose moieties in the glucomannan chain, thus allowing hydrolysis to occur. Notably, enzymes specifically targeting mannans of the PCW are rare in Phytophaga beetles. So far they have been identified and characterized only in *G. viridula* and *Callosobruchus maculatus* (GH5 subfamily 10 or GH5_10) (Pauchet, et al. 2010; Busch, et al. 2017), and one GH5_8 has been characterized in the coffee berry borer *H. hampei* (Acuna, et al. 2012). But in contrast to the activity on glucomannan of some GH45s we observed here, those GH5_10s and GH5_8 were true mannanases, displaying activity towards galactomannan as well as glucomannan. Although our experiments suggested some GH45 cellulases were also active on glucomannan, we believe that the activity these proteins carry out could be important for the degradation of the PCW in the beetle gut. In fact, mannans, including glucomannan, can make up to 5 % of the plant primary cell wall (Scheller and Ulvskov 2010) and may be a crucial enzymatic target during PCW degradation. This hypothesis is further supported by the presence of at least one GH45 protein with some ability to degrade glucomannan in each of the Chrysomelid beetles for which we have functional data.

Another interesting discovery was that several GH45 proteins have lost their ability to use amorphous cellulose as a substrate and evolved instead to degrade xyloglucan, the major hemicellulose of the plant primary cell wall (Pauly, et al. 2013). We believe that the initial substrate shift from cellulose to xyloglucan has likely been promoted by similarities between the substrate backbones (in both cases β-1,4 linked glucose units). The major difference between cellulose and xyloglucan is that the backbone of the latter is decorated with xylose units (which in turn can be substituted by galactose and/or fucose). We presume that the substrate shift from a straight chain polysaccharide such as cellulose to a more complex one such as xyloglucan requires the similar complex adaptation of the enzyme to its novel substrate. However, in contrast to glucomannan-degrading GH45s, GH45 xyloglucanases have apparently completely lost their ability to use amorphous cellulose as a substrate. Here, we clearly demonstrated that, following several rounds of duplications, GH45s in Chrysomelid beetles have evolved novel functions in addition to their ability to break down amorphous cellulose, allowing these insects to degrade two additional major components of the PCW, namely. glucomannan and xyloglucan. This broadening of their functions further emphasizes that GH45 proteins may have likely been an important innovation during the evolution of the Phytophaga beetles and may have strongly contributed to their radiation. In summary, the ability of GH45 proteins to degrade a variety of substrates either as monospecific or as bi-functionalized enzymes indicates that these proteins are particularly prone to substrate shifts.

According to our data, the ability to break down xyloglucan using a GH45 protein has evolved at least twice independently in Phytophaga beetles, once in the LCA of Chrysomelinae and Galerucinae and once in the LCA of the Curculionidae or of the Curculionidae and Brentidae. Once the first Brentidae-derived GH45s are functionally characterized, we will know more. Given that genome/transcriptome data for a majority of families and subfamilies are lacking throughout the Phytophaga clade, we expect that other examples of independent evolution of GH45 xyloglucanases will be revealed in the future. It is important to note that the ability to degrade xyloglucan, which represents an important evolutionary innovation for Phytophaga beetles, may not be linked solely to the evolution of the GH45 family. In fact, in *A. glabripennis* (Cerambycidae: Lamiinae), a glycoside hydrolase family 5 subfamily 2 (GH5_2) protein has evolved to degrade xyloglucan; additionally, orthologous sequences of this GH5_2 xyloglucanase have been found in other species of Lamiinae (McKenna, et al. 2016).

The ability of GH45s to break down xyloglucan correlated with a substitution event from an aspartate to a glutamate residue at a stabilizing site (Asp114) within the Chrysomelidae. Interestingly, the same amino acid exchange was present in SOR3-SO5 but was located at the catalytic acid (Asp121) rather than the stabilizing site (Asp114). Aspartate and glutamate share the same functional group but differ in the length of their side chain. Thus, the preservation of the functional group coupled with an elongated side chain has likely contributed to the substrate switch of those GH45 proteins which when turned on allows xyloglucan to be degraded. Notably, DVI6 and LDE5 do not share that particular substitution but are able to degrade xyloglucan. Therefore, we believe that the transition from cellulase to xyloglucanase has not been driven solely by a single amino acid substitution, but has been triggered by changes at other positions.

Linked to these observations, our Phytophaga-focused phylogeny combined xyloglucanases of Chrysomelidae and Curculionoidea in a single well-supported clade (grouping clades m and n on Fig. 4), suggesting that the LCA of the Phytophaga may have already possessed a GH45 xyloglucanase. Despite the well-supported GH45 xyloglucanase clade, we remain skeptical about a common ancestral GH45 xyloglucanase present in the LCA of the Phytophaga based on two pieces of evidence: first, there are no GH45 xyloglucanases in cerambycid beetles, indicating that Cerambycidae have either lost their GH45 xyloglucanases or never obtained it in the first place. Given that Cerambycidae -- at least species of Lamiinae -- have evolved to degrade xyloglucan by using GH5_2, we believe that a substrate shift from cellulose to xyloglucan of a GH45 never evolved in this family of beetles, not that is was lost. Second, as described above, there are distinct substitution events in the catalytic site between proteins from species of the two superfamilies that likely led to their substrate shift. Based on these facts, we suggest that GH45 xyloglucanases have evolved convergently in both superfamilies. In contrast to GH45 xyloglucanases, GH45 cellulases were present in species of each Phytophaga family investigated to date. This strongly suggests that a cellulolytic GH45 was present in the ancestral Phytophaga species.

According to the carbohydrate-active enzyme (CAZy) database (Lombard, et al. 2014), GH45s encompass 385 sequences (as of February 2018) distributed throughout fungi, bacteria and Metazoans. Interestingly, the distribution of GH45s within Metazoans is rather patchy, encompassing to date only Nematoda (Kikuchi, et al. 2004; Palomares-Rius, et al. 2014), Arthropoda (Pauchet, et al. 2010; Faddeeva-Vakhrusheva, et al. 2016), Rotifera (Szydlowski, et al. 2015) and Mollusca (Sakamoto and Toyohara 2009), and, in insects, is restricted to Phytophaga beetles. We searched several other arthropod genome/transcriptome datasets, including beetles other than Phytophaga, and all publicly available insect genomes, as well as publicly available genomes of Collembola and Oribatida mites. Except for the latter two, we were unable to retrieve GH45 sequences from arthropods. Surprisingly, our phylogenetic analyses clearly showed that the arthropod-derived GH45s, rather than clustering together, formed three separate monophyletic groups. In fact, all metazoan-derived GH45s clustered separately, forming independent monophyletic groups. The patchy distribution of GH45 sequences among Metazoa indicates either that these proteins were acquired multiple times throughout animal evolution or that massive differential gene loss occurred within multicellular organisms. The latter hypothesis appears to be less parsimonious as it implies the existence of multiple GH45s in the LCA of Ophistokonta (Fungi and Metazoa) followed by reciprocal differential gene losses and multiple independent total gene losses in many animal lineages. Intriguingly, the closest clade to the Phytophaga GH45 sequences contained fungal-derived sequences. Our phylogenetic analyses could not identify a specific donor species/group but both suggested species of Saccharomycetales or Neocallimastigaceae fungi as potential source. The most parsimonious explanation for the appearance of GH45 genes in Phytophaga beetles is that one or more genes was acquired by horizontal gene transfer (HGT) from a fungal donor. A similar scenario may have been responsible for the presence of GH45 genes in Oribatida and Collembola, but this hypothesis remains speculative until more sequences from both these orders are identified. In addition to the monophyly of Phytophaga-derived GH45 sequences, a common origin was further suggested by the fact that the position and the phase of the first intron was (except for two cases) conserved across GH45 genes from the species of Cerambycidae, Chrysomelidae and Curculionidae for which genome data are available. If our hypotheses are correct, the LCA of all Phytophaga beetles most likely acquired a single GH45 gene from a fungal donor. As we did not find any fungal (Saccharomycetales or Neocalimastigaceae) introns corresponding to the proposed original intron, it appears that beetle-derived GH45 genes have acquired an intron after the HGT. The GH45 gene then likely underwent several duplications before the separation of the different Phytophaga clades, and these duplications continued independently after the diversification of this hyper-diverse clade of beetles.

Strikingly, nematode-derived GH45 sequences were consistently grouped together with Saccharomycetales fungi in each analysis we ran, clearly demonstrating that the closest relatives to their GH45 genes were fungal and from different fungi than the insect relatives. The origin of nematode-derived GH45 genes has been investigated, and their acquisition by HGT from a fungal source has been proposed (Kikuchi, et al. 2004; Palomares-Rius, et al. 2014). Here we provide the third independent confirmation of this fact.

In conclusion, our research indicated that the Phytophaga GH45s have adapted to substrate shifts. In addition to cellulose, this adaptation led to the recognition and catalysis of two additional substrates, neither of which can be enzymatically addressed by any other GH family that those insects encode. Beetles of the Chrysomelidae have evolved to break down three components of the PCW (cellulose, xyloglucan and glucomannan) by using only GH45s. In concert with GH28 pectinases encoded by each investigated species (Kirsch, et al. 2014), these beetles have evolved a near-complete set of enzymatic tools with which to deconstruct the PCW, allowing them to gain access to the nutrient-rich plant cell contents; in addition, PCW-derived polysaccharides are a potential source of energy. Our data also suggest that GH45 is not an ancestral gene family but was likely acquired by the LCA of the Phytophaga from a fungal source, most likely through an HGT event. Both, enzymatic activity and ancestral origin suggest that GH45s were likely an essential prerequisite for the adaptation allowing Phytophaga beetles to feed on plants.

## Materials and methods

### Production of recombinant GH45 proteins

Open reading frames (ORFs) were amplified from cDNAs using gene-specific primers based on previously described GH45 sequences of *C. tremula, P. cochleariae, L. decemlineata, D. virgifera virgifera* and *S. oryzae* (Pauchet, et al. 2010). If necessary, full-length transcript sequences were obtained by rapid amplification of cDNA ends PCR (RACE-PCR) using RACE-ready cDNAs as described by (Pauchet, et al. 2010). For downstream heterologous expression, ORFs were amplified using a forward primer designed to include a Kozak sequence and a reverse primer designed to omit the stop codon. cDNAs initially generated for the (RACE-PCR), as described by (Pauchet, et al. 2010), were used as PCR template, and the PCR reactions were performed using a high-fidelity Taq polymerase (AccuPrime, Invitrogen). The PCR products were cloned into the pIB/V5-His TOPO (Invitrogen) in frame with a V5-(His)_6_ epitope. TOP10 chemically competent *Escherichia coli* cells (Invitrogen) were transformed and incubated overnight on a LB agar plate containing ampicillin (100 μg/ml). To select constructs for which the recombinant DNA had ligated in the correct orientation, randomly picked colonies were checked by colony PCR using the OpIE2 forward primer (located on the vector) and a gene-specific reverse primer (Table S5). Positive clones were further cultured in 2x yeast extract tryptone (2xYT) medium containing 100 μg/ml ampicillin. After plasmid isolation using GeneJET Plasmid Miniprep Kit (Thermo Scientific), the recombinant plasmids were sequenced in both directions using Sanger sequencing to confirm whether the ORF has been correctly inserted into the vector. Positive constructs were then transfected in *Sf*9 insect cells (Invitrogen) using FuGENE HD (Promega) as a transfection reagent. First, successful expression was determined by transiently transfecting three clones per construct in a 24-well plate format. 72 h after transfection, the culture medium was harvested and centrifuged (16,000 x g, 5 min, 4 °C) to remove cell debris. Successful expression was verified by Western blot using the anti-V5-HRP antibody (Invitrogen). In order to collect enough material for downstream enzymatic activity assays, a single clone per construct was chosen to be transiently transfected in a 6-well plate format. 72 h after transfection, culture medium was harvested and treated as described above. The cell medium was stored at 4 °C until further use.

### Enzymatic characterization

The enzymatic activity of recombinant proteins was initially tested on agarose diffusion assays using carboxymethyl cellulose (CMC) as a substrate. Agarose (1%) plates were prepared, containing 0.1 % CMC in 20 mM citrate/phosphate buffer pH 5.0. Small holes were made in the agarose matrix using cut-off pipette tips, to which 10 μl of the crude culture medium of each expressed enzyme was applied. After incubation for 16 h at 40 °C, activity was revealed by incubating the agarose plate in 0.1% Congo red for 1 h at room temperature followed by washing with 1 M NaCl until pale halos on a red background were visible. To investigate GH45 enzymatic activity in more detail, we analyzed their enzymatic breakdown products using thin layer chromatography (TLC). For that, the culture medium of transiently transfected cells was dialyzed and desalted as described in Busch et al. (2017). The following substrates were tested: CMC, Avicel, glucomannan, galactomannan and xyloglucan (all from Megazyme) with a final concentration of 0.5 %. We also tested regenerated amorphous cellulose (RAC), prepared according to Zhang et al. (2006). Additionally, we used the cello-oligomers D-(+)-biose to D-(+)-hexaose (all from Megazyme), as substrates at a final concentration of 250 ng/μl. Samples were incubated and analyzed as previously described (Busch, et al. 2017). The reference standard contained 2 μg of each oligomer: glucose, cellobiose, cellotriose, cellotetraose and cellopentaose as well as isoprimeverose, xylosyl-cellobiose and the hepto-, octa- and nona saccharides of xyloglucan.

### Tissue-specific gene expression

Third-instar *P. cochleariae* larvae, actively feeding on leaves of *B. rapa*, were used for total RNA extraction. Larvae were cut open from abdomen to head, and the complete gut was removed and stored separately from the rest of the body. Dissection and storage were carried out in RL solution (Analytik Jena). Three biological replicates were sampled, each containing three larvae. Total RNA was isolated using the innuPREP RNA Mini Kit (Analytik Jena), following the manufacturer’s protocol. The resulting RNA samples were subjected to DNase digestion (Ambion), and their quality was assessed using the RNA 6000 Nano LabChip kit on a 2100 Bioanalyser (both Agilent Technologies). Total RNA was used as a template to synthesize cDNAs, using the Verso cDNA synthesis kit (Thermo Scientific). The resulting cDNA samples were then used for real-time quantitative PCR (qPCR) experiments, which were performed in 96-well hard-shell PCR plates on the CFX Connect Real-Time System (both Bio-Rad). All reactions were carried out using the 2-Step QPCR SYBR Kit (Thermo Scientific), following the manufacturer’s instructions. Primers were designed using the program Primer3 (version 0.4.0) (Table S5). The specific amplification of each transcript was verified by dissociation curve analysis. A standard curve for each primer pair was determined in the CFX Manager (version 3.1) based on Cq-values (quantitation cycle) of qPCRs run with a dilution series of cDNA pools. The efficiency and amplification factors of each qPCR, based on the slope of the standard curve, were calculated using an integrated efficiency calculator of the CFX manager software (version 3.1). Ribosomal protein S3 (RPS3), extracted from our *P. cochleariae* larval gut transcriptome (Kirsch, et al. 2012), was used as a reference gene, and the abundance of GH45 transcripts was expressed as RNA molecules per 1000 RNA molecules of RPS3. Gene expression values were ln-transformed and significant differences between gut and rest-body were analyzed in SigmaPlot Version 11.0 using paired t-tests.

### Gene structure determination

Genomic sequences of GH45 encoding genes were mined from publicly available draft genomes of *L. decemlineata* (Schoville, et al. 2018), *A. glabripennis* (McKenna, et al. 2016), *H. hampei* (Vega, et al. 2015), *D. ponderosae* (Keeling, et al. 2013) and *S. oryzae* (unpublished; accession: SAMN08382431). The intron/exon structure was determined for each gene using splign (Kapustin, et al. 2008), a spliced aligner.

### Amino acid alignment and Phytophaga-specific phylogenies

Sequences corresponding to Phytophaga GH45 proteins described in our previous studies were combined with those mined from several NCBI databases, such as the non-redundant protein database (ncbi_nr) and the transcriptome shotgun assembly database (ncbi_tsa) (Table S4). In addition, transcriptome datasets generated from species of Phytophaga beetles were retrieved from the short-read archive (ncbi_sra) (Table S4) and assembled using the CLC workbench program version 11.0. Reads were loaded and quality trimmed before being assembled using standard parameters. The resulting assemblies were screened for contigs matching known beetle GH45 sequences through BLAST searches. The resulting contigs were then manually curated and used for further analysis. Amino acid alignments were carried out using MUSCLE version 3.7 implemented in MEGA7 (version 7.0.26) (Kumar, et al. 2016). The maximum likelihood analysis was also conducted in MEGA7. The best model of protein evolution was determined in MEGA7 using the ‘find best DNA/protein models’ tool. The best model was the ‘Whelan and Goldman’ (WAG) model, incorporating a discrete gamma distribution (shape parameter = 5) to model evolutionary rate differences among sites (+G) and a proportion of invariable sites (+I). The robustness of the analysis was tested using 1,000 bootstrap replicates.

### Large phylogenetic analysis

We used the GH45 protein sequence from *Sitophilus oryzae* (ADU33247.1) as a BLASTp query against the NCBI’s non-redundant protein library with an E-value threshold of 1E^−3^. We retrieved the 250 best blast hits (Table S3), encompassing a majority of fungal sequences as well as various hexapod sequences (including Chrysomelidae, Curculionidae, Lamiinae and Collembola (=Entomobryomorpha)). Besides fungi and hexapods, GH45 sequences from 10 nematodes, from one Tardigrade, one Rotifer and one bacterium, as well as a few uncharacterized protists from environmental samples, were among the 250 best BLAST hits. We complemented this dataset with predicted proteins from several Oribatid mites (10 sequences) and Collembola (4 sequences) retrieved from ncbi_tsa (Table S3). This resulted in a collection of 264 sequences.

The set of 264 protein sequences was scanned against the Pfam v31 library of protein domains using the pfam_scan script with default parameters. All these proteins had one GH45 (Glyco_hydro_45 PF02015) domain on at least 98% of the expected lengths. Most of the fungal proteins possessed an ancillary carbohydrate-binding module (either CBM1 or CBM10), but this domain was not found in other species, except the bdelloid rotifer *Adineta ricciae*. We eliminated redundancy at 90% identity level between the 264 protein sequences; we used the CD-HIT Suite server (Huang, et al. 2010) and reduced the dataset to 201 non-redundant sequences, while maintaining the diversity of clades.

The non-redundant sequences were aligned using MAFFT v7.271 (Katoh and Standley 2013) with the “—auto” option to allow us to automatically select the most appropriate alignment strategy. We used trimal (Capella-Gutierrez, et al. 2009) to automatically discard columns that contained more than 50% of gaps in the alignment (-gt 0.5 option) and the maximum likelihood and the Bayesian methods to reconstruct phylogenetic trees. Maximum likelihood trees were reconstructed by RAxML version 8.2.9 (Stamatakis 2014) with an estimated gamma distribution of rates of evolution across sites and an automatic selection of the fittest evolutionary model (PROTGAMMAAUTO). Bootstrap replicates were automatically stopped upon convergence (-autoMRE). Bayesian trees were reconstructed by MrBayes version 3.2.6 (Ronquist, et al. 2012)with an automatic estimation of the gamma distribution of rates of evolution across sites and a mixture of evolutionary models. The number of mcmc generations was stopped once the average deviation of split frequencies was below 0.05. Twenty-five percent of the trees were burnt for calculation of the consensus tree and statistics.

## Supporting information

## Acknowledgments

We are grateful to Bianca Wurlitzer and Domenica Schnabelrauch for technical support. We thank Emily Wheeler, Boston, for editorial assistance. We express our gratitude to Franziska Beran (MPI for Chemical Ecology) for sharing the transcriptomes of *Phyllotreta armoraciae* and *Psylliodes chrysocephala* prior to publication. We are also thankful to Roy Kirsch and David G. Heckel for their input on experimental design and for fruitful discussions. This work was supported by the Max Planck Society.

---

**Table S1.**
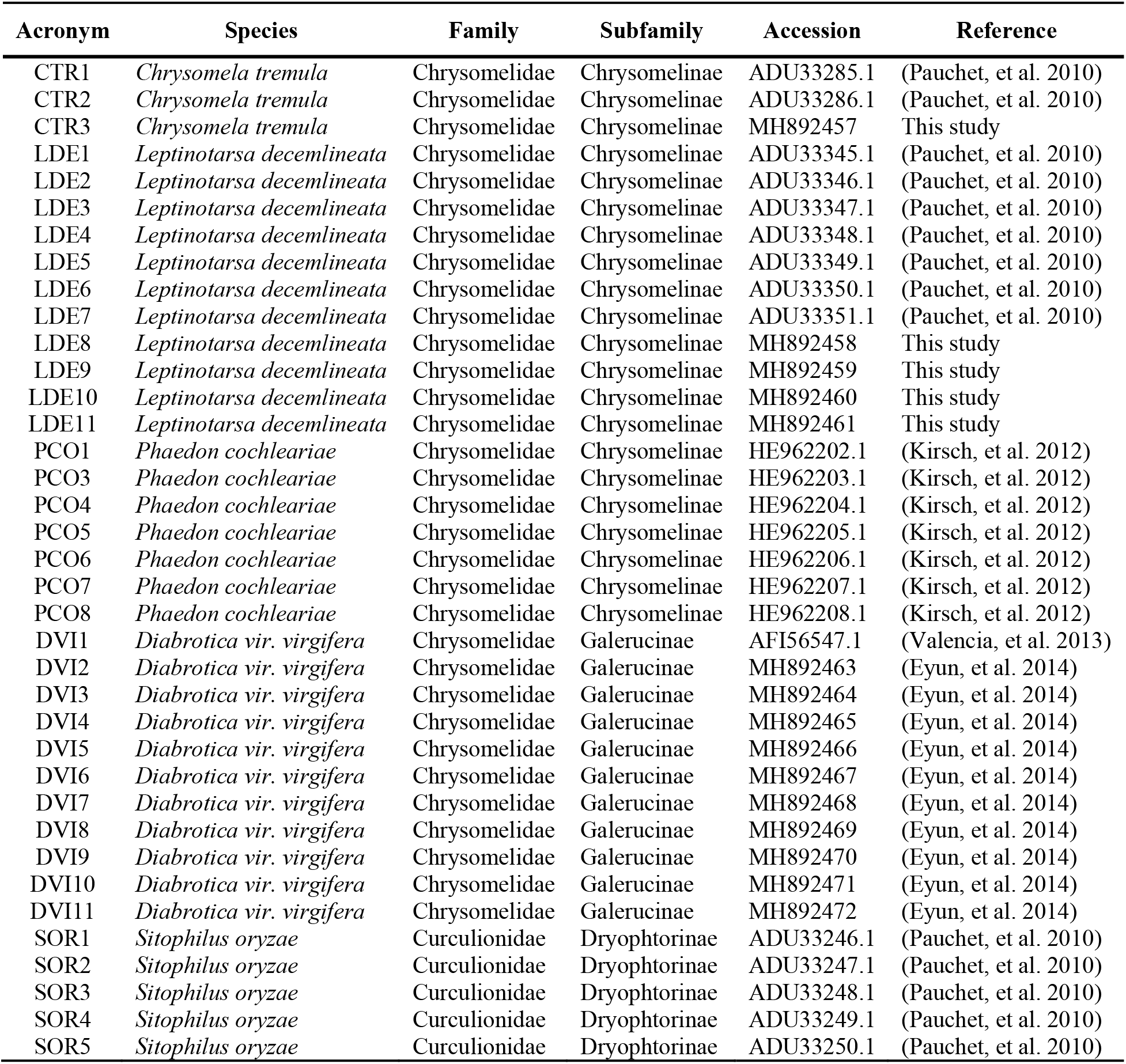
Details on the beetle-derived GH45 proteins that were expressed in *Sf*9 cells.

**Table S2.** Significant differences of genes expressed in different tissues

|  | t-value | p-value |
| --- | --- | --- |
| Pco1 | 14.04 | 0.005 |
| Pco3 | 8.732 | 0.013 |
| Pco4 | 6.706 | 0.022 |
| Pco5 | 7.004 | 0.02 |
| Pco6 | 5.913 | 0.027 |
| Pco7 | 5.695 | 0.029 |
| Pco8 | 34.908 | <0.001 |

**Table S3.**
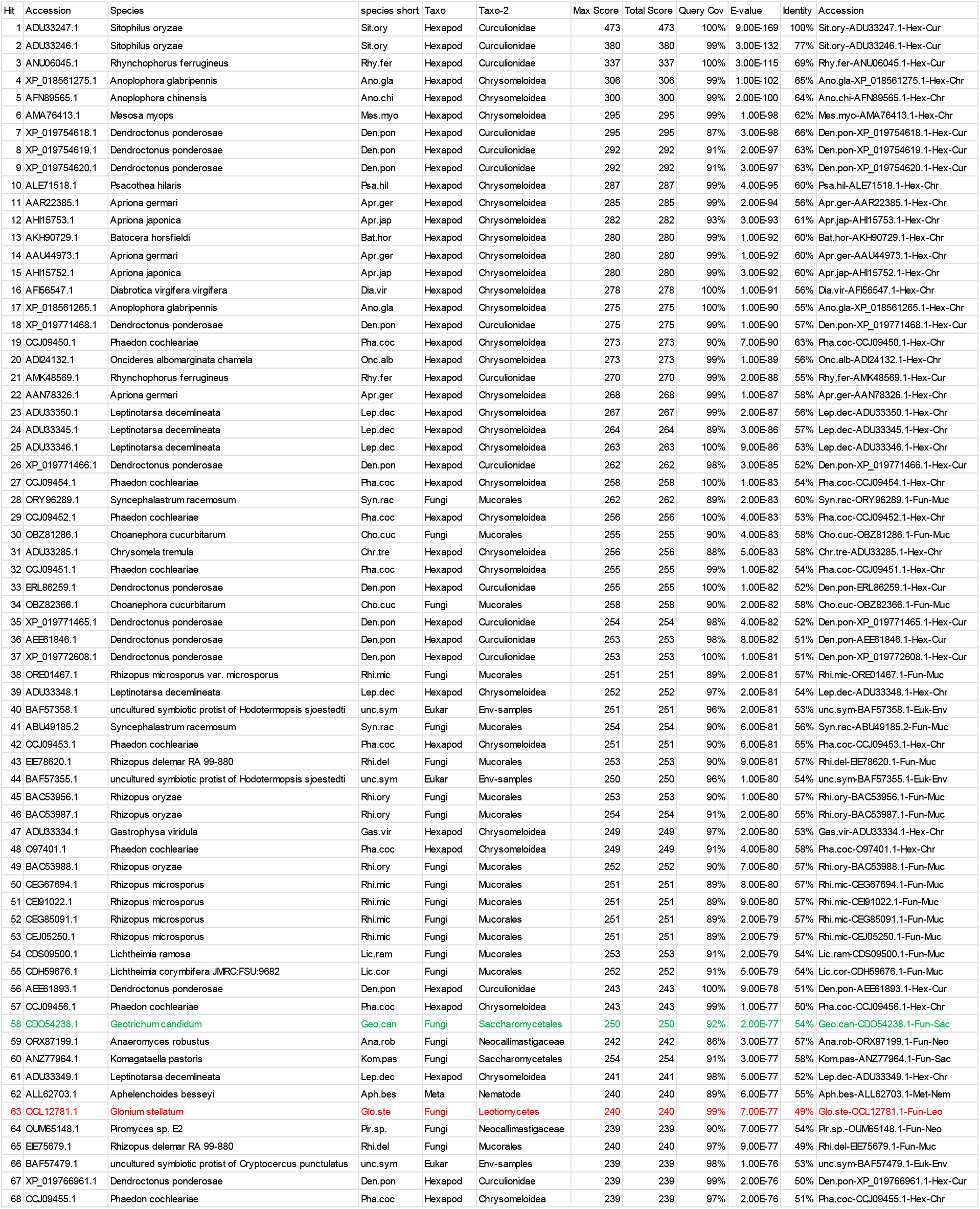
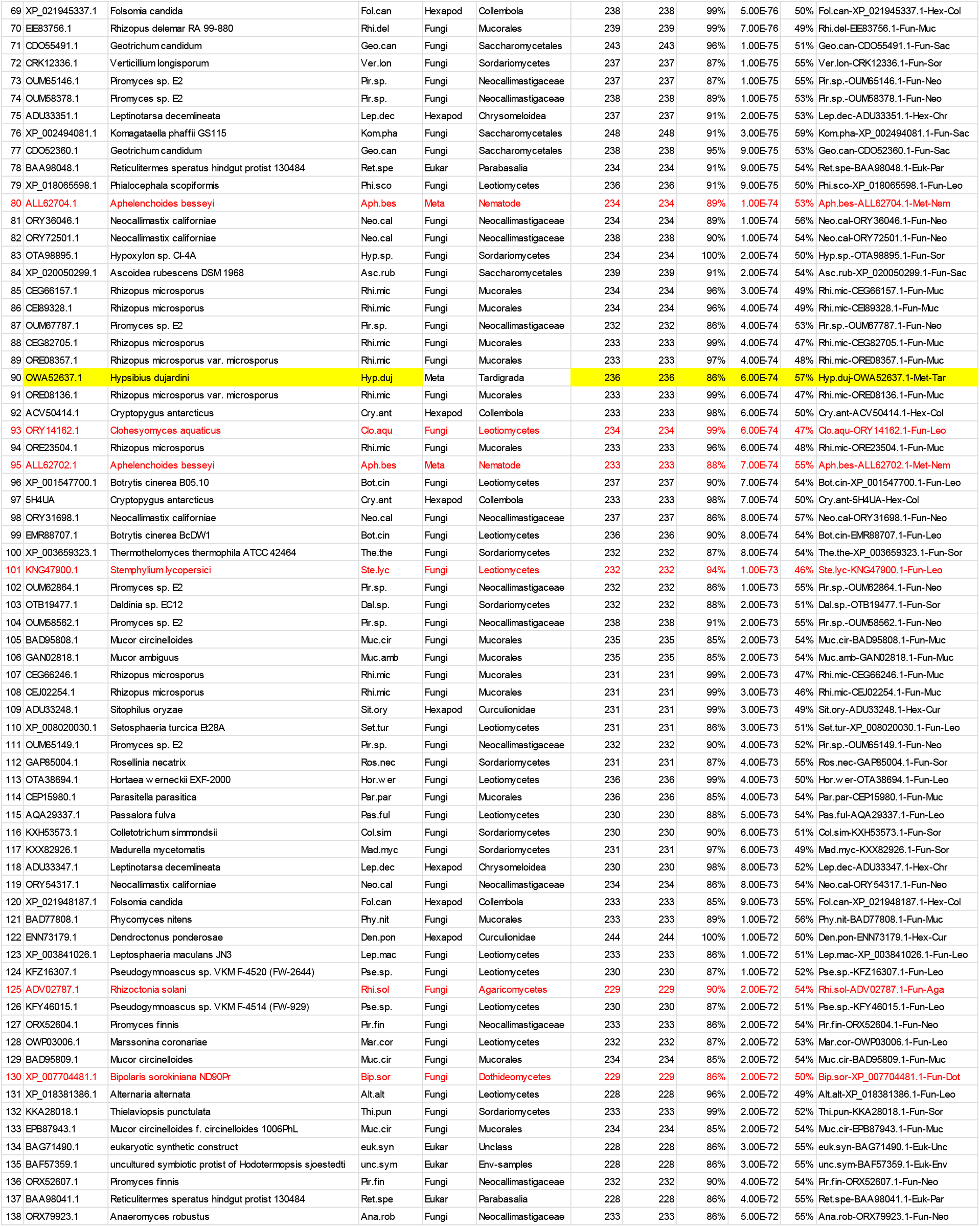
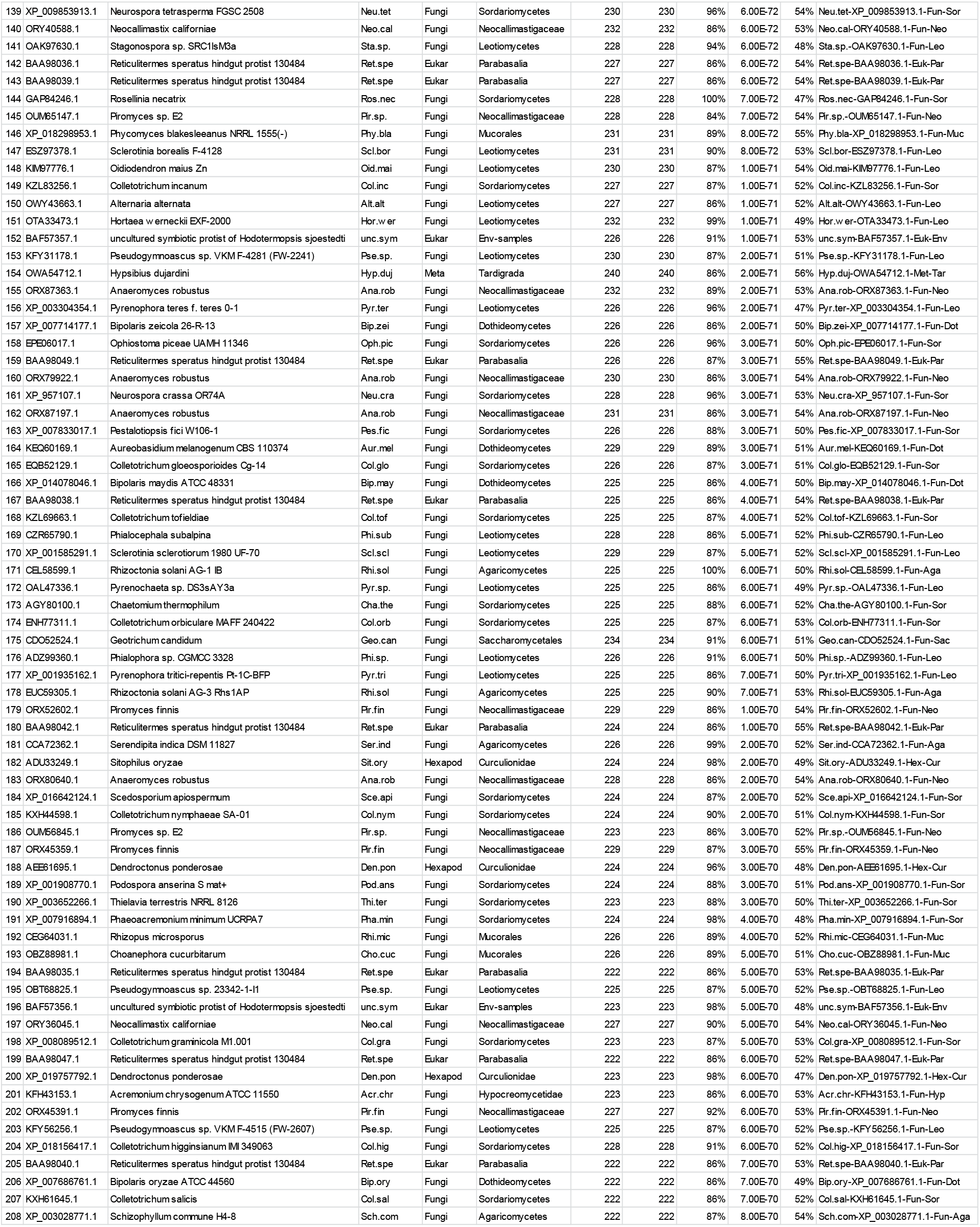
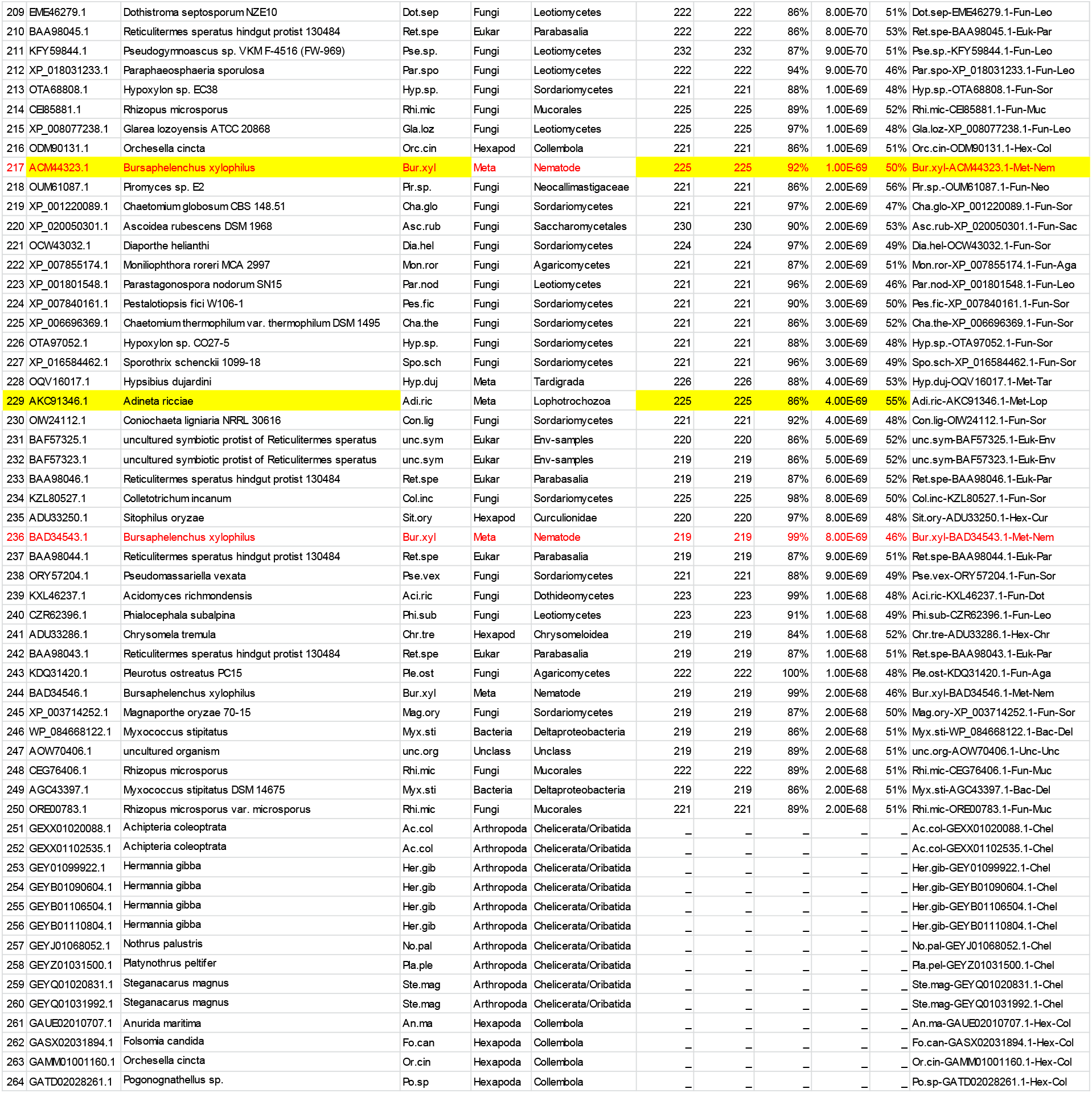
Details of the sequences used for the large phylogenetic analyses.

**Table S4.**
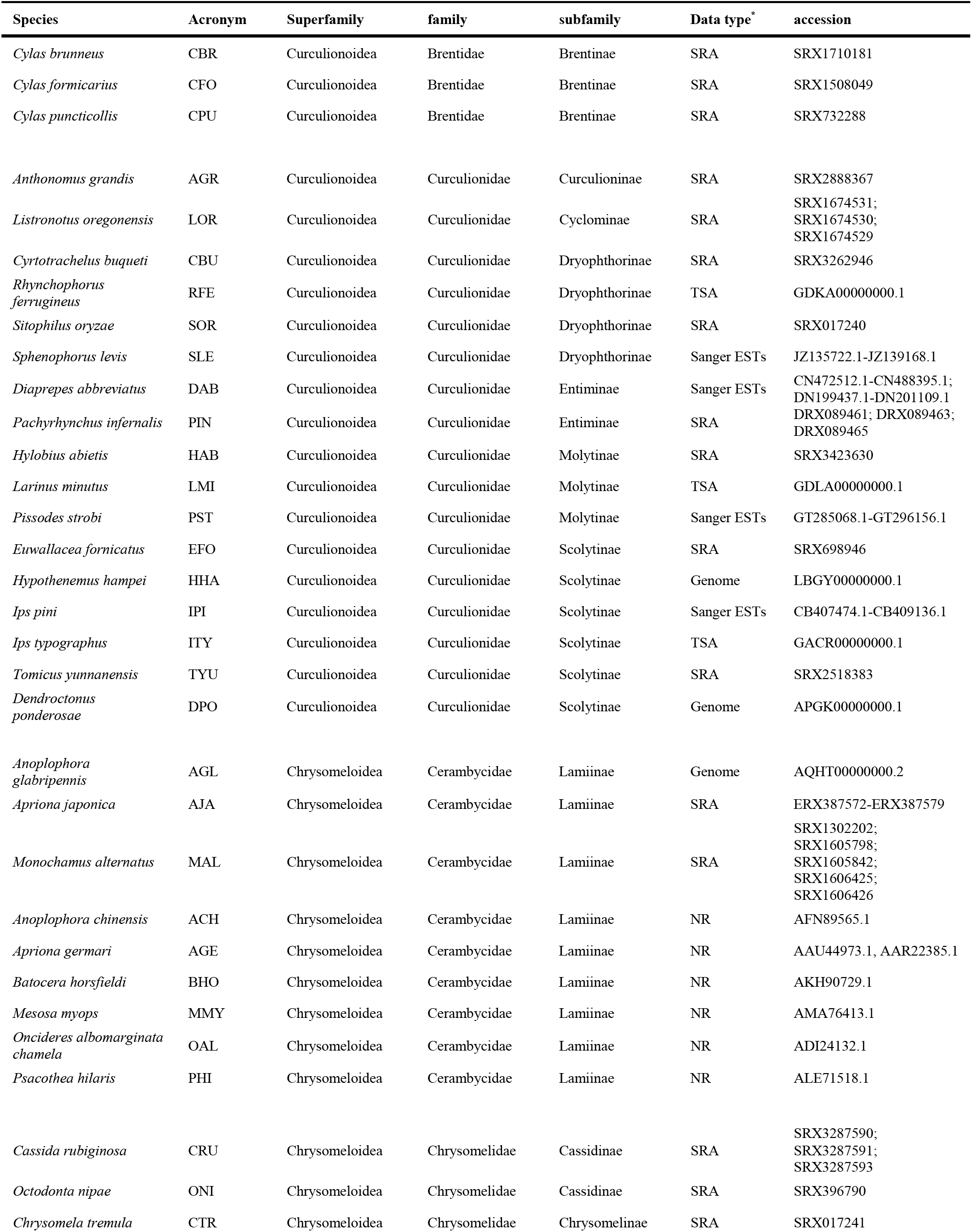
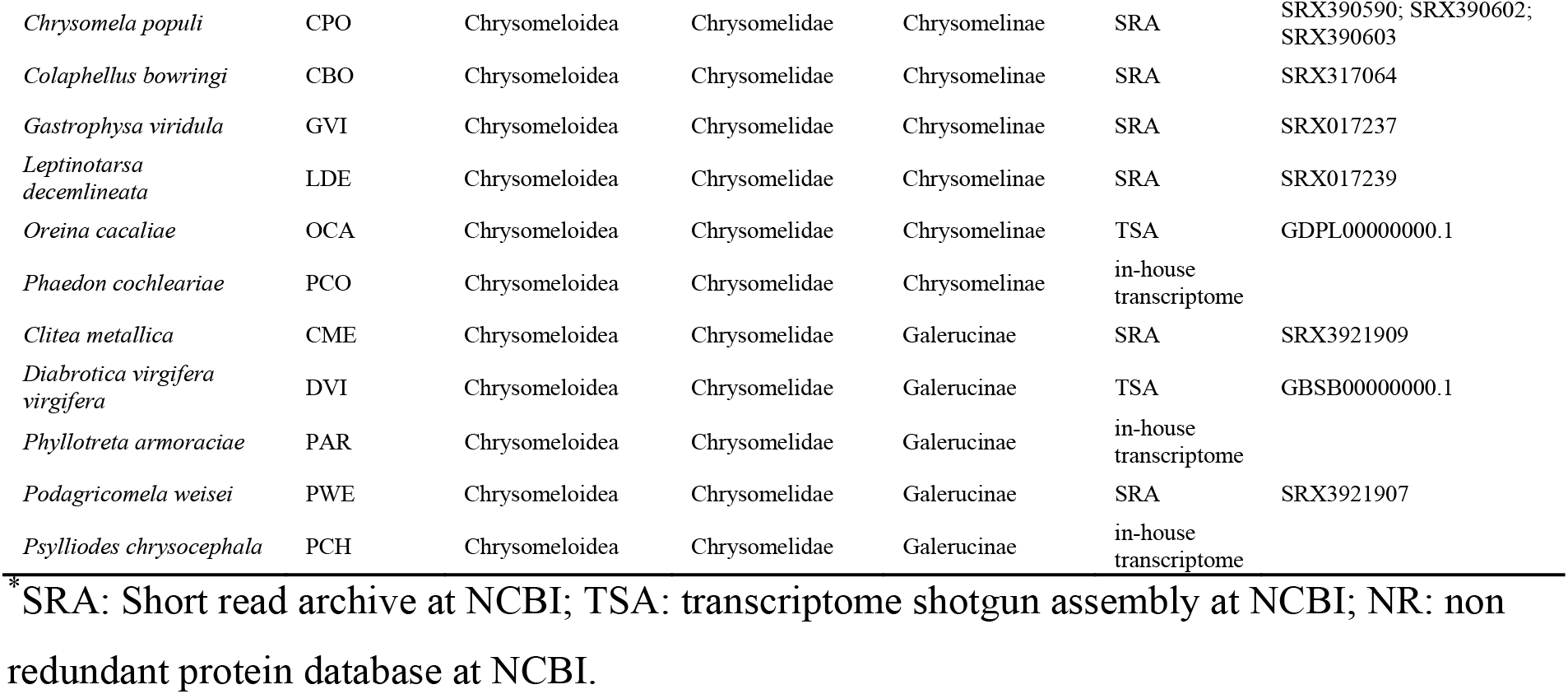
Details on the genome/transcriptome datasets which were used to curate GH45 sequences derived from Phytophaga beetles.

**Table S5.**
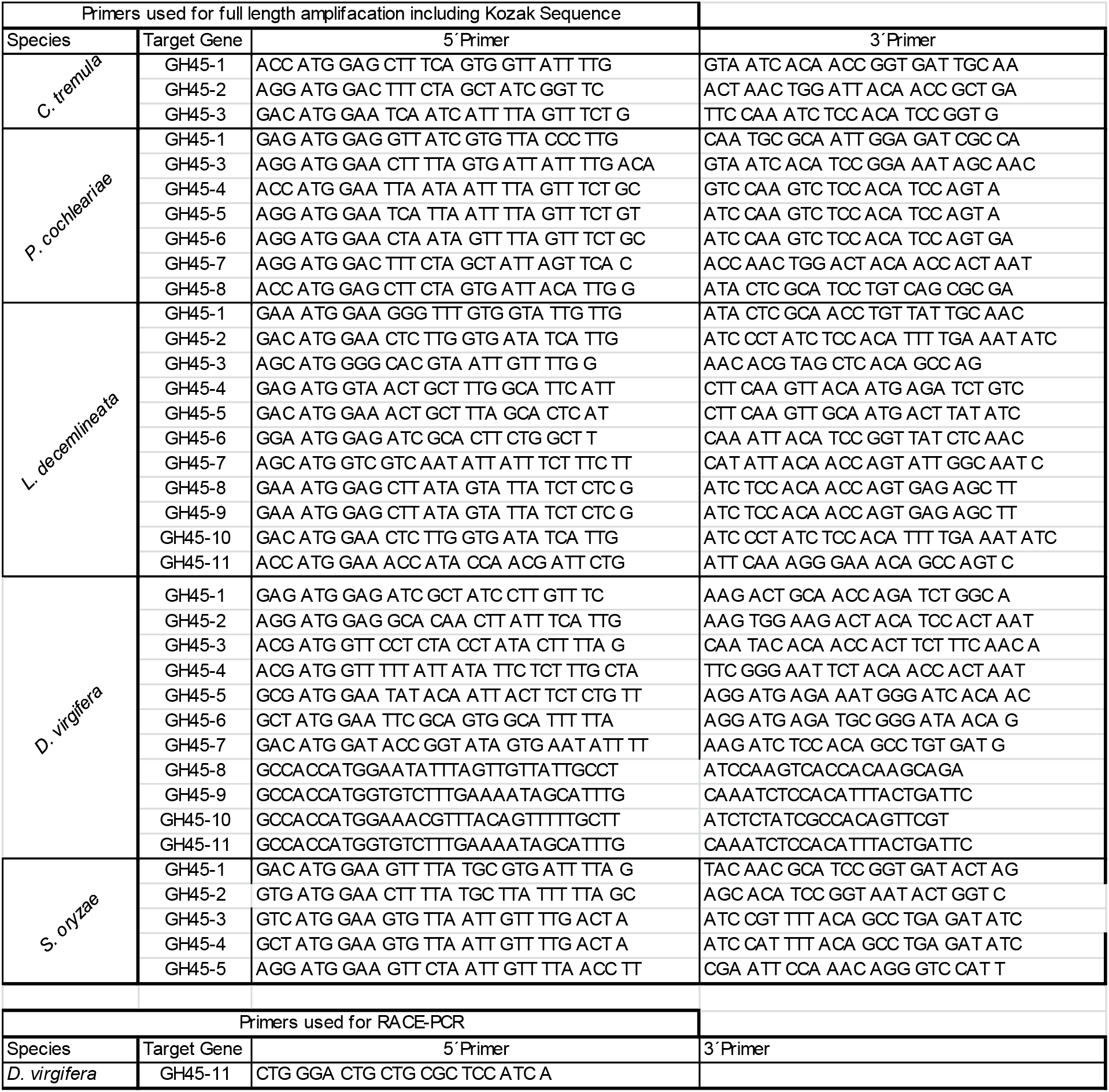
List of primers used in this study

## Supplementary Figure Legends

**Fig. S1 Thin-layer chromatography of *C. tremula* GH45s assayed against several plant cell wall polysaccharides**. Recombinant GH45s were incubated for 16 h at 40 °C with various plant polysaccharides. Their breakdown products were analyzed on TLC and visualized using 0.2 % orcinol in methane/sulphoric acid (9:1) under continuous heating. Each TLC represents an individually tested GH45 (Ctr1 to Ctr3). All GH45s were assayed against the same set of substrates, namely, cellotriose to cellohexaose (C3-C6); crystalline cellulose = avicel (CC); carboxymethyl cellulose (CMC); regenerated amorphous cellulose (RAC); glucomannan (GluM); galactomannan (GalM); xyloglucan (XG); standards: C1 = glucose, C2 – C5 = cellobiose to pentaose; XG2-XG9 = xyloglucan-oligomers.

**Fig. S2 Thin layer chromatography of *P. cochleariae* GH45s assayed against several plant cell wall polysaccharides**. Recombinant GH45s were incubated for 16 h at 40 °C with various plantpolysaccharides. Their breakdown products were analyzed on TLC and visualized using 0.2 % orcinol in methane/sulphoric acid (9:1) under continuous heating. Each TLC represents an individually tested GH45 (Pco1 to Pco8). All GH45s were assayed against the same set of substrates which included cellotriose to cellohexaose (C3-C6); crystalline cellulose = avicel (CC); carboxymethyl cellulose (CMC); regenerated amorphous cellulose (RAC); glucomannan (GluM); galactomannan (GalM); xyloglucan (XG); standards: C1 = glucose, C2 – C5 = cellobiose – pentaose; XG2-XG9 = xyloglucan-oligomers.

**Fig. S3 Thin layer chromatography of *L. decemlineata* GH45s assayed against several plant cell wall polysaccharides**. Recombinant GH45s were incubated for 16 h at 40 °C with various plant polysaccharides. Their breakdown products were analyzed on TLC and visualized using 0.2 % orcinol in methane/sulphoric acid (9:1) under continuous heating. Each TLC represents an individually tested GH45 (Lde1 to Lde11). All GH45s were assayed against the same set of substrates which included: cellotriose to cellohexaose (C3-C6); crystalline cellulose = avicel (CC); carboxymethyl cellulose (CMC); regenerated amorphous cellulose (RAC); glucomannan (GluM); galactomannan (GalM); xyloglucan (XG); standards: C1 = slucose, C2 – C5 = cellobiose – pentaose; XG2-XG9 = xyloglucan-oligomers.

**Fig. S4 Thin-layer chromatography of *D. virgifera* GH45s assayed against several plant cell wall polysaccharides**. Recombinant GH45s were incubated for 16 h at 40 °C with various plant polysaccharides. Their breakdown products were analyzed on TLC and visualized using 0.2 % orcinol in methane/sulphoric acid (9:1) under continuous heating. Each TLC represents an individually tested GH45 (Dvi1 to Dvi11). All GH45s were assayed against the same set of substrates: cellotriose to cellohexaose (C3-C6); crystalline cellulose = avicel (CC); carboxymethyl cellulose (CMC); regenerated amorphous cellulose (RAC); glucomannan (GluM); galactomannan (GalM); xyloglucan (XG); standards: C1 = Glucose, C2 – C5 = cellobiose – pentaose; XG2-XG9 = xyloglucan-oligomers.

**Fig. S5 Thin-layer chromatography of *S. oryzae* GH45s assayed against several plant cell wall polysaccharides**. Recombinant GH45s were incubated for 16 h at 40 °C with various plant polysaccharides. Their breakdown products were analyzed on TLC and visualized using 0.2 % orcinol in methane/sulphoric acid (9:1) under continuous heating. Each TLC represents an individually tested GH45 (Sor1 toSor5). All GH45s were assayed against the same set of substrates: cellotriose to cellohexaose (C3-C6); crystalline cellulose = avicel (CC); carboxymethyl cellulose (CMC); regenerated amorphous cellulose (RAC); glucomannan (GluM); galactomannan (GalM); xyloglucan (XG); standards: C1 = glucose, C2 – C5 = cellobiose – pentaose; XG2-XG9 = xyloglucan-oligomers.

**Fig. S6 Bayesian global phylogenetic analysis encompassing GH45 proteins from various taxa (expanded version of Figure 4)**. 264 GH45 sequences of microbial and metazoan origin were initially collected (see Methods) and their redundancy was eliminated at 90 % sequence similarity, resulting in a total of 201 sequences. Sequence details are given in Table S3. Fungal branches are marked in orange, symbiotic protists in red, Collembola, Oribatida and Entognatha in dark blue, Coleoptera in light blue, Nematoda and Tardigrada in dark green, Rotifera in light green and bacteria in purple.

**Fig. S7 Maximum likelihood inferred phylogenetic analysis encompassing GH45 proteins from various taxa**. 264 GH45 sequences of microbial and metazoan origin were initially collected (see Methods), and their redundancy was eliminated at 90 % sequence similarity, resulting in a total of 201 sequences. Sequence details are given in Table S3. Fungal branches are marked in orange, symbiotic protists in red, Oribatida in dark blue, Collembola, Entognatha and Coleoptera in light blue, Nematoda in light green, Tardigrada and Rotifera in dark green and bacteria in purple.

**Fig. S8 Conservation of intron position in phytophagous beetles with known GH45 genome structure**. Amino acid alignment of GH45 sequences derived from four different phytophagous beetles using MUSCLE. Genomic sequence information was retrieved from the genome assemblies of *L. decemlineata* (Schoville, et al. 2018), *H. hampei* (Vega, et al. 2015), *A. glabripennis* (McKenna, et al. 2016) and *D. ponderosae* (Keeling, et al. 2013). The predicted signal peptide is marked in bold letters. Intron positions are highlighted by colored amino acids according to their phase. Phase 0: green; Phase 1: red; Phase 2: blue.

**Fig. S9 Amino acid alignment of the GH45 catalytic residues based on our Curculionoidea-based phylogeny**. We used a GH45 sequence of Humicola insulens (HIN1) as a reference sequence (Accession: 2ENG_A) (Davies, et al. 1995). According to HIN1, we chose to investigate the catalytic residues (ASP10 and ASP121) as well as a conserved tyrosine (TYR8) of the catalytic binding site, a crucial substrate stabilizing amino acid (ASP114) and an essential conserved alanine (ALA74). Arrows indicate amino acid residue under investigation. If highlighted in green, the residue remained unchanged in comparison to HIN1; elsewise it is highlighted in red. GH45 enzymatic activity was color-coded based on the respective substrate specificity (green dots = endo-β-1,4-glucanase, blue dots = endo-β-1,4-xyloglucanase, red dots= no activity). Color coding in reference to the respective subfamily: pink = Scolytinae (Curculionidae); brown = Entiminae (Curculionidae); purple = Cyclominae (Curculionidae); gray = Curculioninae (Curculionidae); yellow = Molytinae (Curculionidae); light blue = Brentinae (Brentidae); dark blue = Dryophthorinae (Curculionidae). Each clade corresponds to the clades depicted in Fig. 4.

**Fig. S10 Amino acid alignment of the GH45 catalytic residues using our Chrysomeloidea-based phylogeny**. We used a GH45 sequence of Humicola insulens (HIN1) as a reference sequence (Accession: 2ENG_A) (Davies, et al. 1995). According to HIN1, we chose to investigate the catalytic residues (ASP10 and ASP121) as well as a conserved tyrosine (TYR8) of the catalytic binding site, a crucial substrate stabilizing amino acid (ASP114) and an essential conserved alanine (ALA74). Arrows indicate amino acid residue under investigation. If highlighted in green, the residue remained unchanged in comparison to HIN1, otherwise it is highlighted in red. GH45 enzymatic activity was color-coded based on the respective substrate specificity (green dots = endo-β-1,4-glucanase, blue dots = endo-β-1,4-xyloglucanase, red dots = no activity). Color-coding was performed with reference to the respective subfamily: dark green = Chrysomelinae (Chrysomelidae); light green = Galerucinae (Chrysomelidae); orange = Lamiinae (Cerambycidae); cyan = Cassidinae (Chrysomelidae). Each clade corresponds to the clades depicted in Fig. 4.

