## Supplementary figures and images for "Functional analyses of the horizontally acquired Phytophaga glycoside hydrolase family 45 (GH45) proteins reveal distinct functional characteristics"

### Supplementary file 1

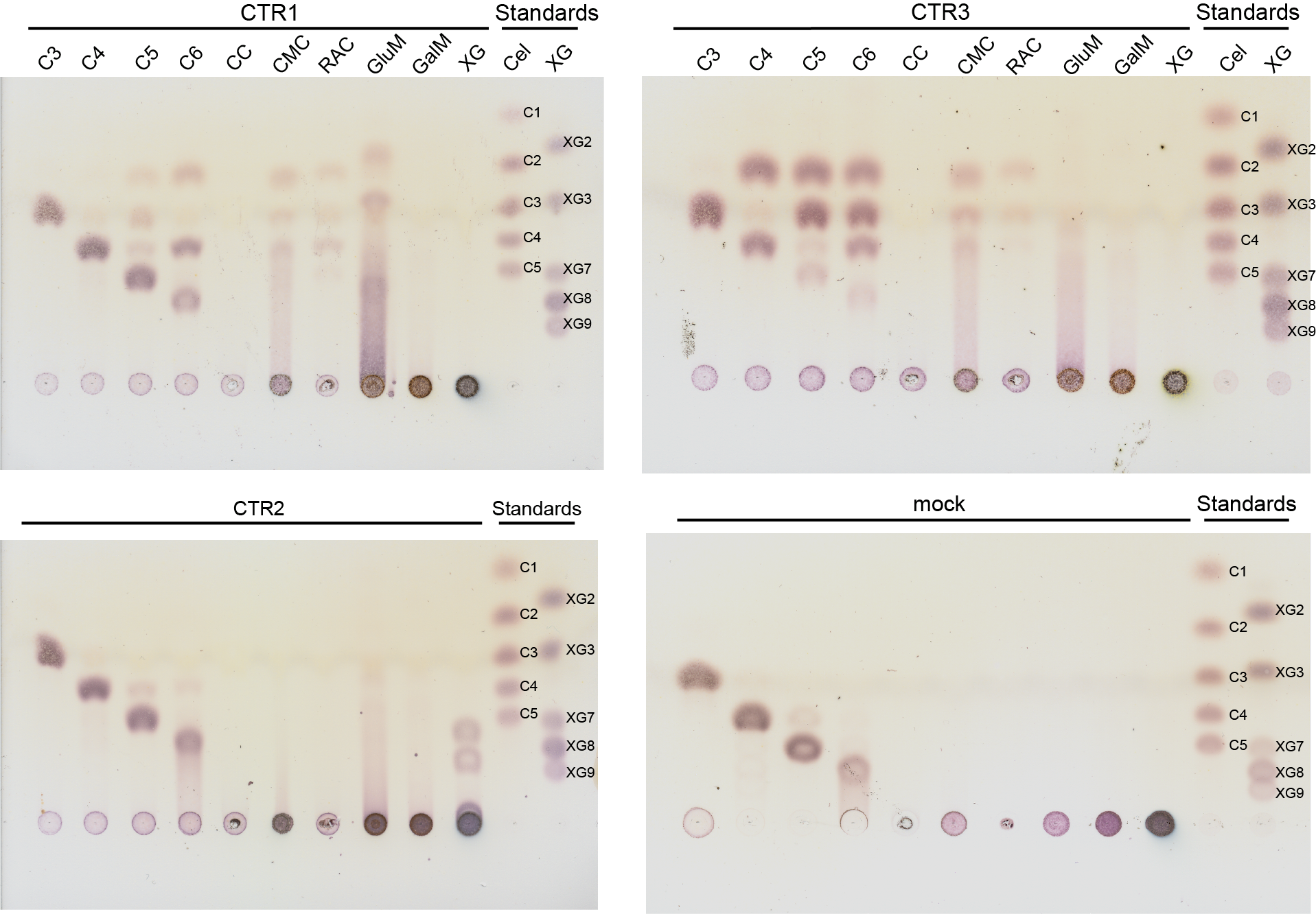

### Supplementary file 2

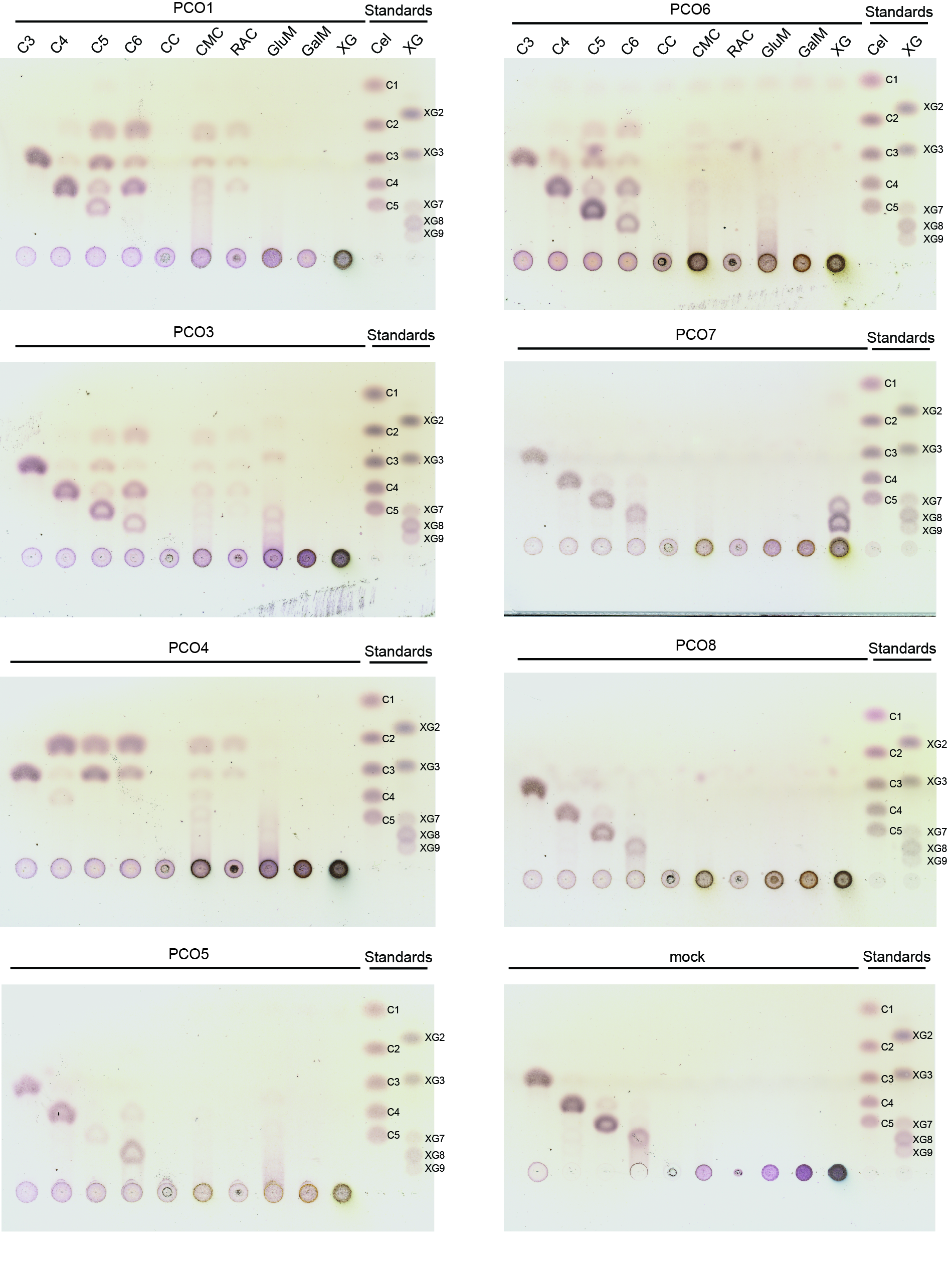

### Supplementary file 3

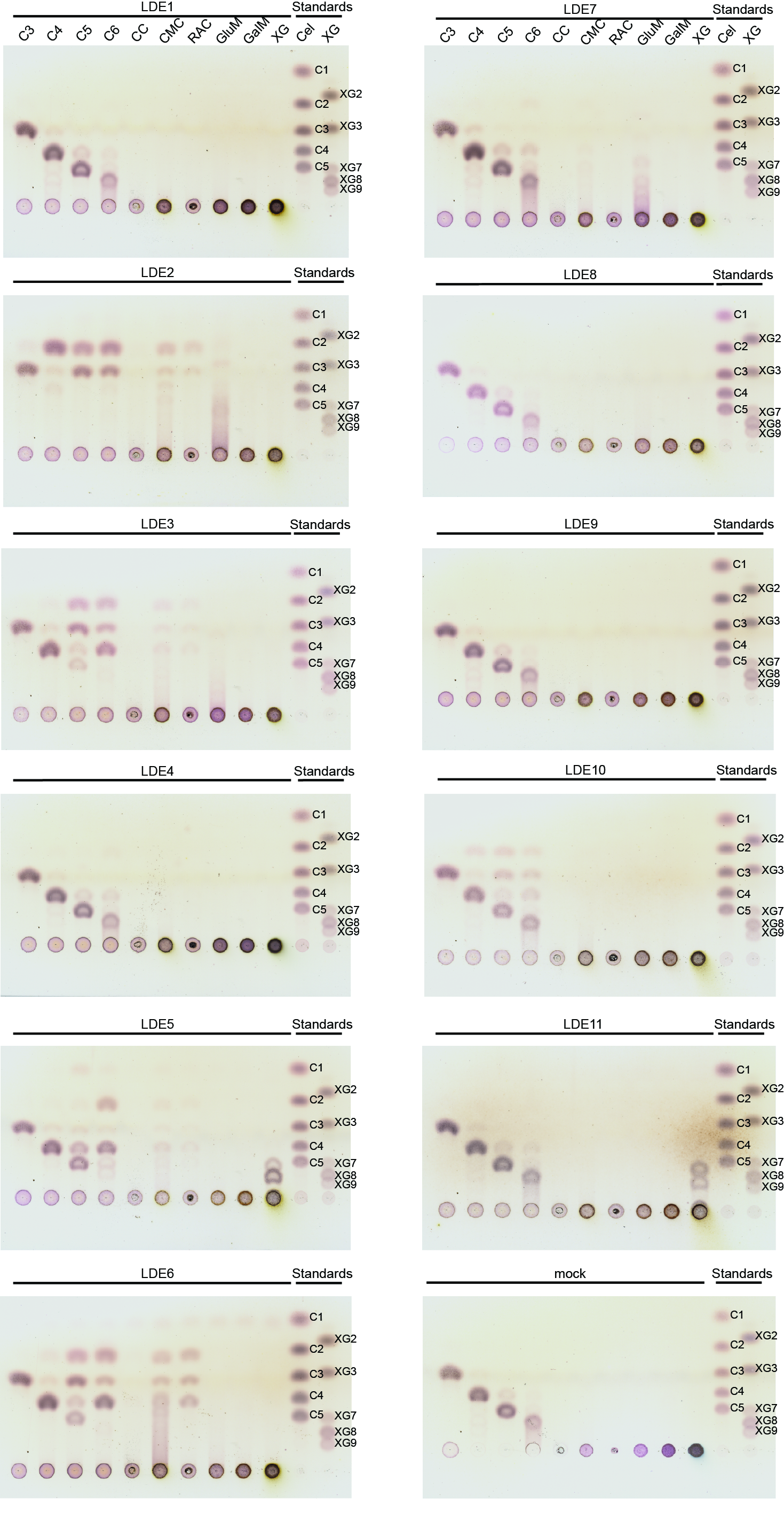

### Supplementary file 4

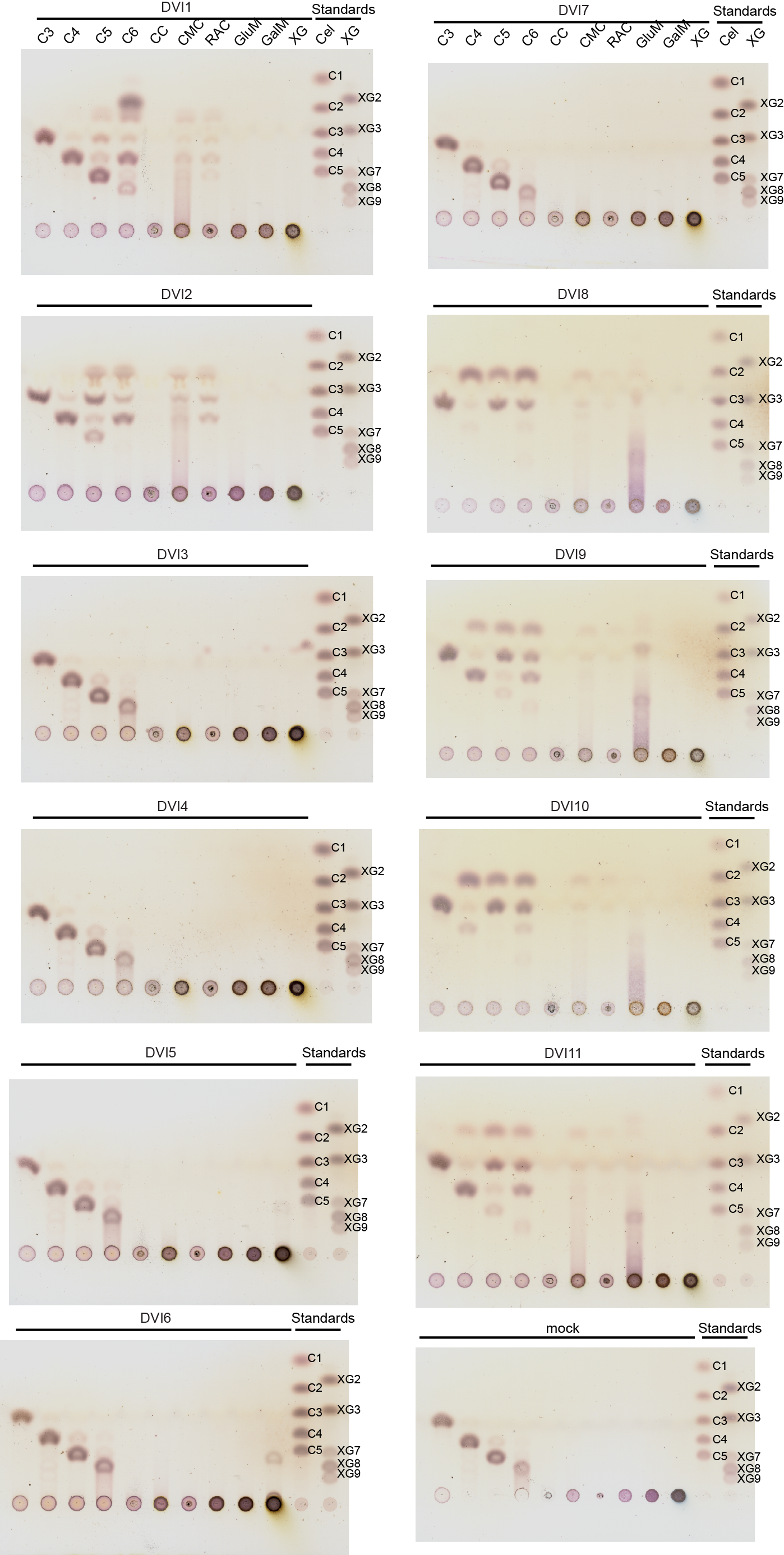

### Supplementary file 5

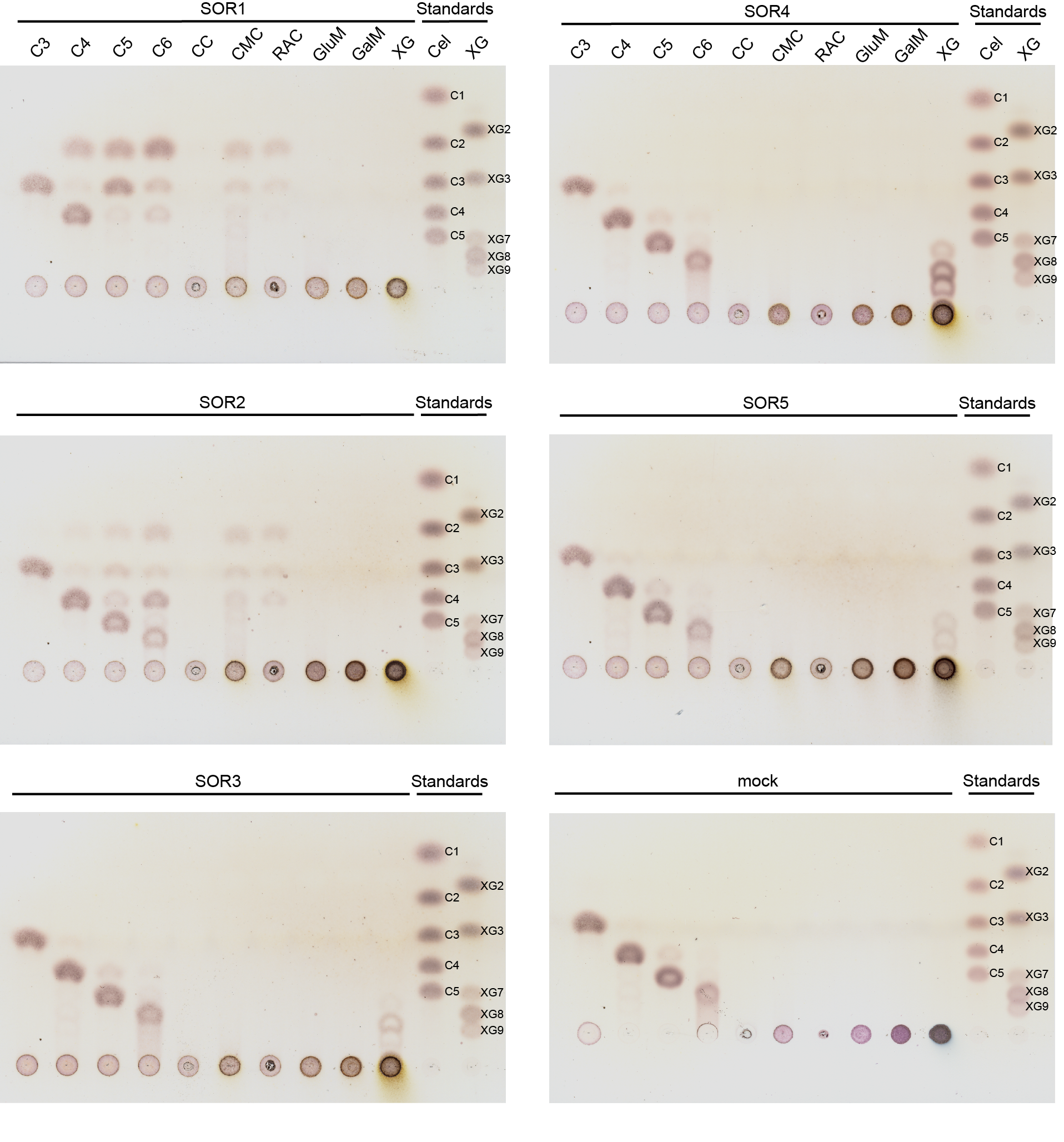

### Supplementary file 6

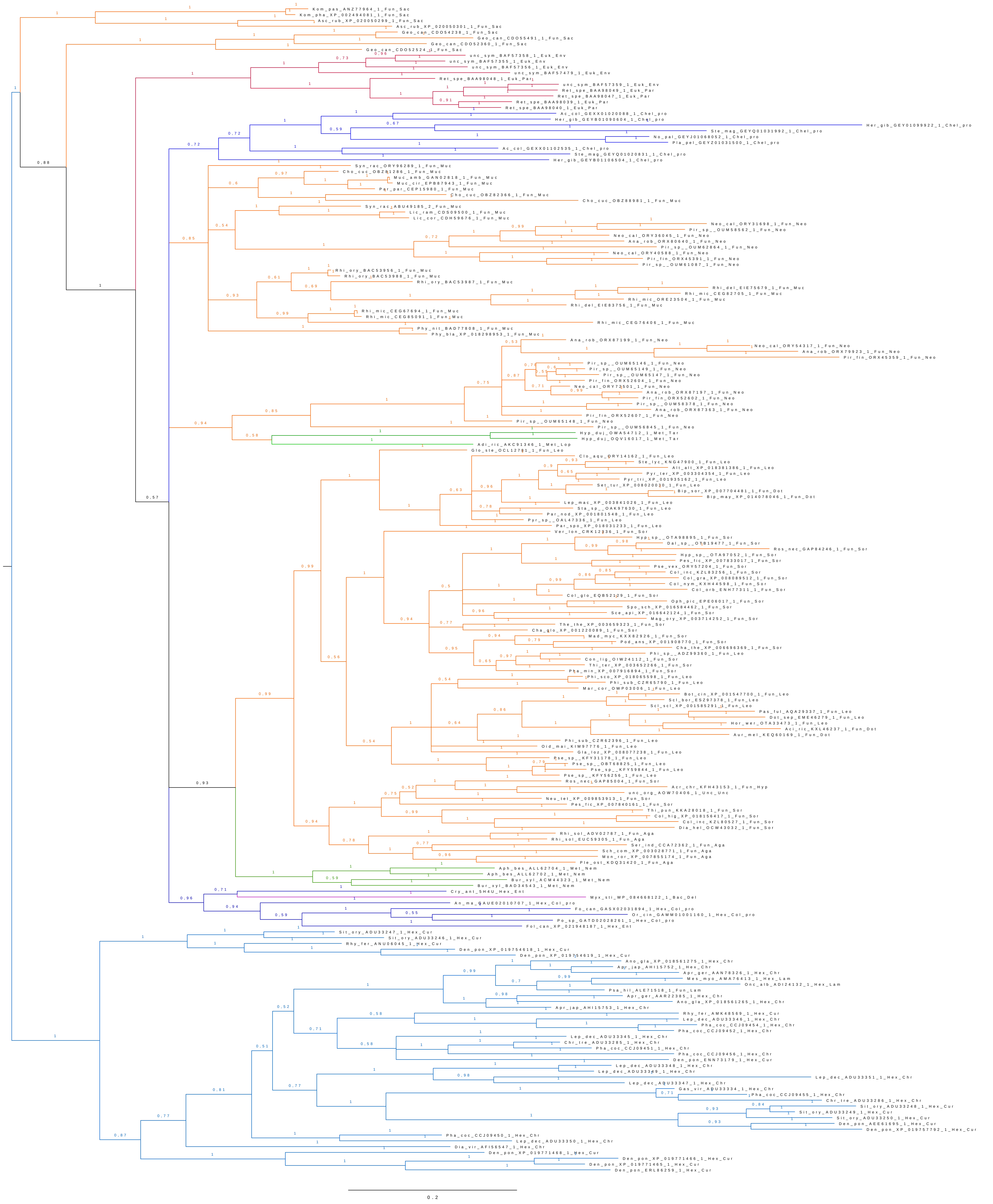

### Supplementary file 7

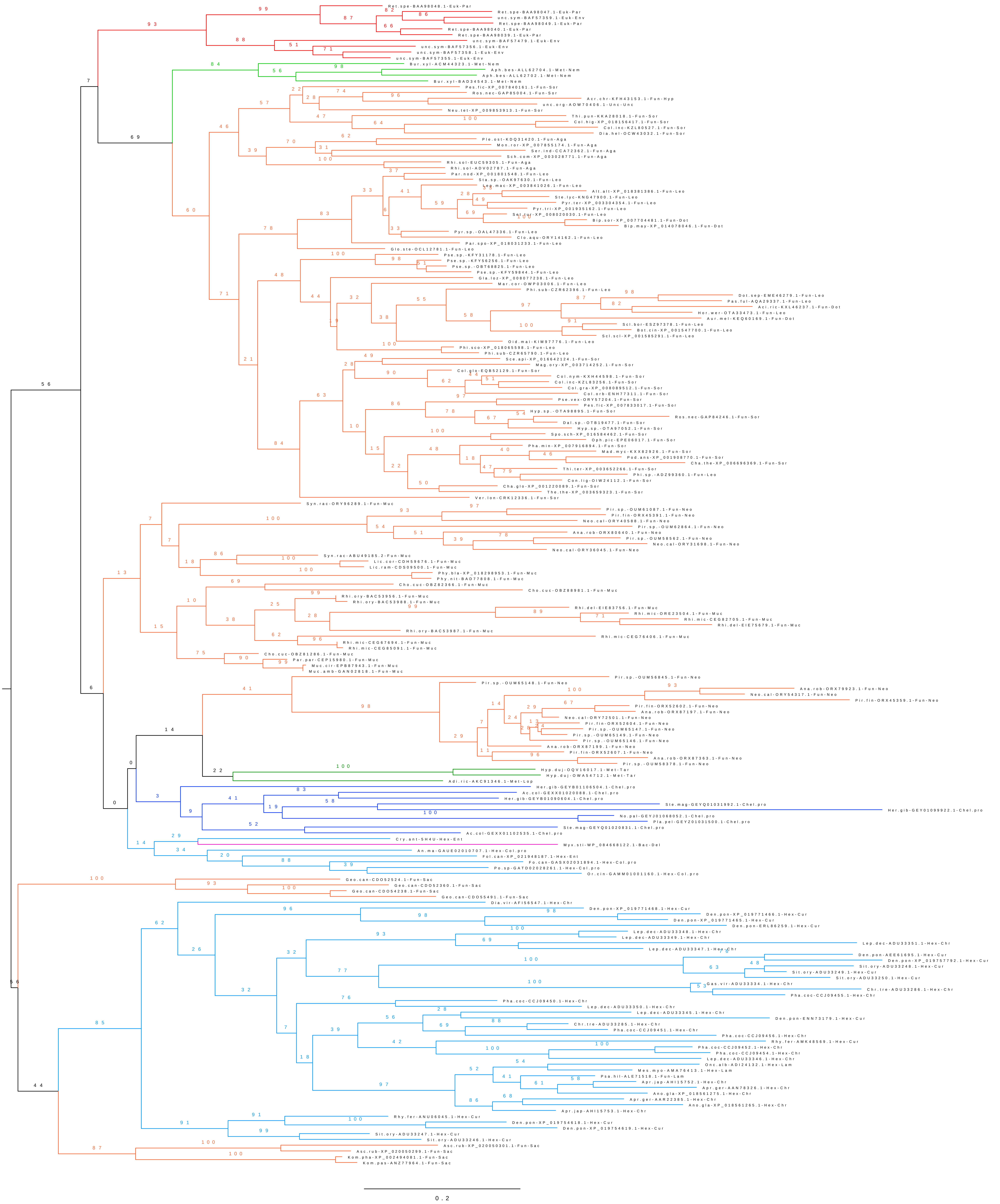

### Supplementary file 8

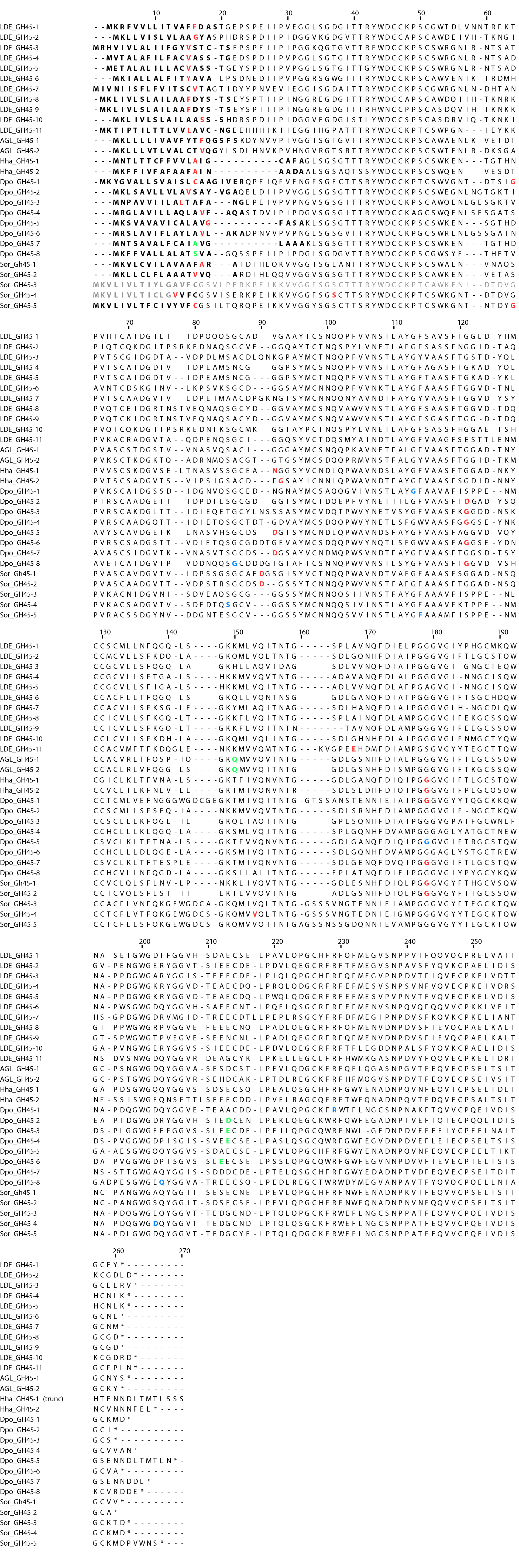

### Supplementary file 9

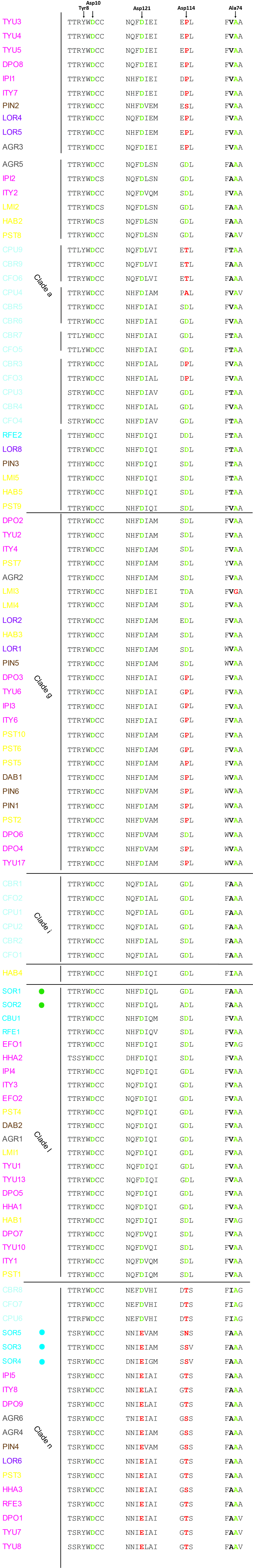

### Supplementary file 10

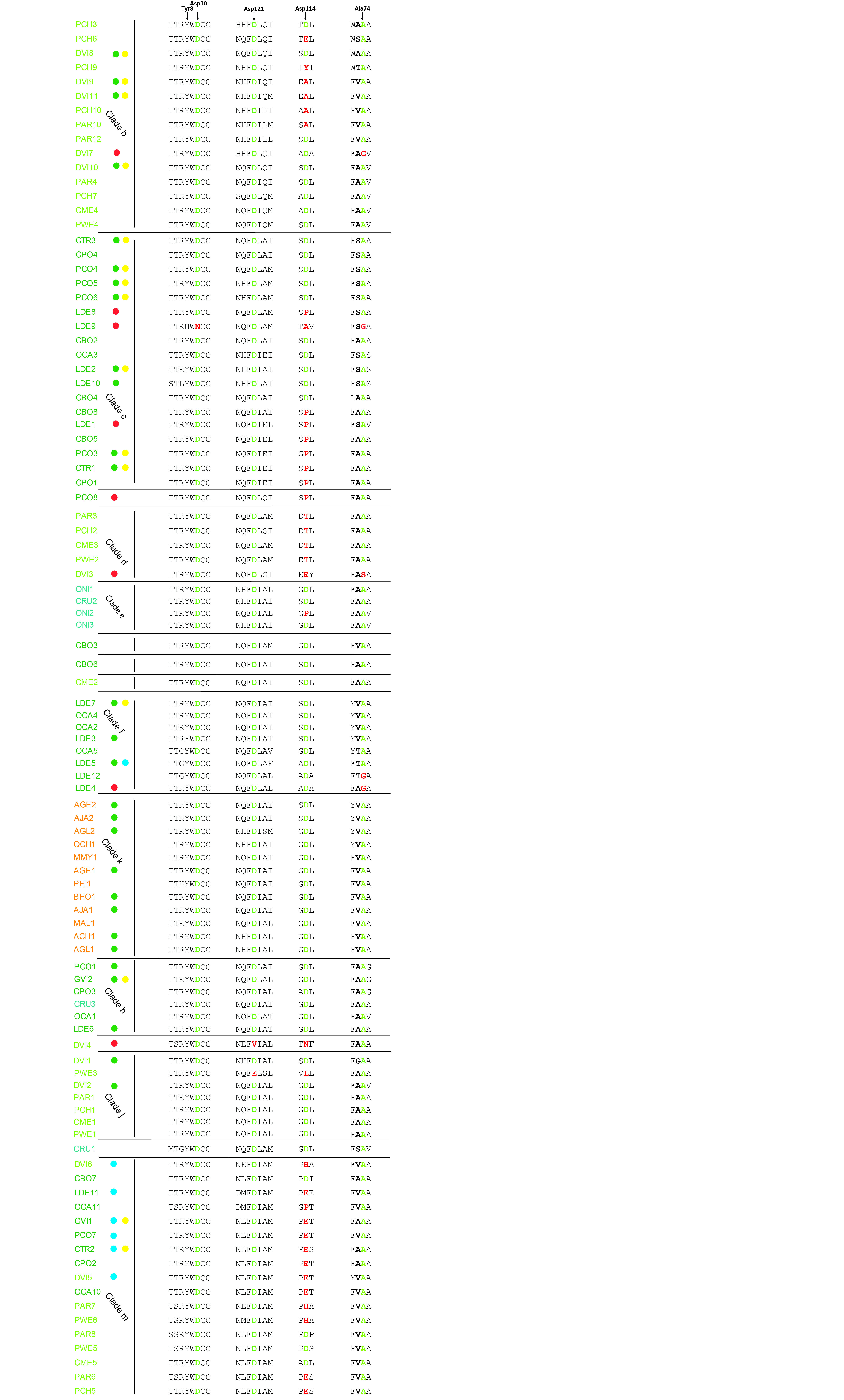
